# N-terminal oligomerization drives HDAC4 nuclear condensation and neurodevelopmental dysfunction in *Drosophila*

**DOI:** 10.1101/2025.03.10.642474

**Authors:** Hannah R Hawley, Andrew J Sutherland-Smith, Matthew S Savoian, Helen L Fitzsimons

## Abstract

Histone deacetylase four (HDAC4) undergoes dynamic nucleocytoplasmic shuttling, which is important in the regulation of its activity. However, aberrant nuclear accumulation of HDAC4 is associated with both neurodevelopmental and neurodegenerative disease, and in our *Drosophila* model, impairs normal neuronal development. Upon nuclear accumulation, HDAC4 forms biomolecular condensates, previously termed aggregates, that correlate with the severity of defects in development of the *Drosophila* mushroom body and adult eye. Here we determined that nuclear condensation of HDAC4 is dependent on self-oligomerization, and that impairing oligomerization reduces condensation and the severity of neurodevelopmental phenotypes in *Drosophila*. HDAC4 condensates are highly dynamic and are stabilized by the presence of MEF2, which promotes their formation and coalescence, ultimately exacerbating phenotypic severity. These data provide insight into the role of HDAC4 condensates in normal neuronal function and suggest that their disruption may contribute to the onset or progression of neuronal disease. Consequently, targeting oligomerization of HDAC4 and its interaction with MEF2 present potential therapeutic strategies for diseases associated with HDAC4 nuclear accumulation.

## Introduction

Histone deacetylase four (HDAC4) is an important regulator of neuronal development and memory formation in species across the animal kingdom including *C. elegans*, *Drosophila*, rodents and humans (Fitzsimons *et al*., 2013; Kim *et al*., 2012; Sando *et al*., 2012; Trazzi *et al*., 2016; Wakeling *et al*., 2021; Wang *et al*., 2011; Zhu *et al*., 2019). HDAC4 is widely expressed in the rodent brain (Broide *et al*., 2007; Darcy *et al*., 2010) and detected in all regions examined in the human protein atlas (Sjöstedt *et al*., 2020). As a member of the Class IIa family of histone deacetylases, it is characterised by a conserved deacetylation domain and an extended N-terminal region that interacts with regulatory proteins including transcription factors (Davis *et al*., 2003; Miska *et al*., 1999; Vega *et al*., 2004; Wang & Yang, 2001). While early studies identified high sequence conservation with catalytically active HDACs (Hassig *et al*., 1998) and deacetylase activity *in vitro* (Grozinger *et al*., 1999), vertebrate HDAC4 harbors little intrinsic deacetylase activity due to an inactivating mutation (Lahm *et al*., 2007), instead facilitating deacetylation via association with HDAC3 and the NCoR/SMRT repressor complex (Fischle *et al*., 2001; Fischle *et al*., 2002; Huang *et al*., 2000). Although HDAC4 interacts with transcription factors and chromatin-associated complexes, it shuttles between the nucleus and cytoplasm, and localizes predominantly to the cytoplasm in many neuronal subtypes (Bolger & Yao, 2005; Chen *et al*., 2014; Darcy *et al*., 2010; Litke *et al*., 2018; Mielcarek *et al*., 2013b; Sando *et al*., 2012; Schlumm *et al*., 2013; Tan *et al*., 2024; Zhu *et al*., 2019). Altered subcellular distribution of HDAC4 has been observed in a number of neurodevelopmental and neurodegenerative diseases, therefore recent efforts have focused on increasing our understanding of the regulation of HDAC4 shuttling, as well as its roles in both the nucleus and cytoplasm.

The distribution of HDAC4 in neurons is governed by its intrinsic nuclear localization and export sequences, as well as via co-factors including MEF2 and 14-3-3 proteins, which mediate nuclear import and export, respectively (Fitzsimons, 2015). As the interaction between HDAC4 and 14-3-3 is phosphorylation-dependent (Grozinger & Schreiber, 2000; Wang *et al*., 2000), nucleocytoplasmic shuttling of HDAC4 is dynamic and occurs in response to synaptic activity in neurons (Chawla *et al*., 2003; Chen *et al*., 2014; Litke *et al*., 2018; Sando *et al*., 2012; Schlumm *et al*., 2013; Zhu *et al*., 2019). Activation of Ca^2+^/calmodulin kinases leads to the phosphorylation of three serine residues (Ser246, Ser467 and Ser632 in human HDAC4), which mediate 14-3-3 binding and promote the nuclear export of HDAC4 (Grozinger & Schreiber, 2000; McKinsey *et al*., 2000; Miska *et al*., 2001; Wang *et al*., 2000). Conversely, dephosphorylation by protein phosphatase 2a (PP2A) results in loss of 14-3-3 binding, reducing the efficiency of nuclear export and cytoplasmic anchoring (Paroni *et al*., 2008), which enables MEF2-dependent nuclear import of HDAC4 (Borghi *et al*., 2001; Wang & Yang, 2001). Given this dynamic regulation, the localization of HDAC4 differs depending on the neuronal subtype in which it is expressed (Darcy *et al*., 2010).

Dysregulation of HDAC4, particularly its aberrant accumulation in the nucleus, has been associated with several neurodegenerative and neurodevelopmental disorders. Increased nuclear concentration of HDAC4 has been observed in the brains of individuals with Alzheimer’s disease (AD) (Herrup *et al*., 2013; Shen *et al*., 2016) and Ataxia Telangiectasia (Li *et al*., 2012), as well as in mouse models of AD (Colussi *et al*., 2023; Sen *et al*., 2015; Shen *et al*., 2016), CDKL5 disorder (Trazzi *et al*., 2016), 2q37 deletion syndrome (Sando *et al*., 2012) and Parkinson’s disease (Lang *et al*., 2019; Wu *et al*., 2016). HDAC4 staining in neurons displays a punctate, granular pattern (Darcy *et al*., 2010; Tan *et al*., 2024), which is likely due to self-oligomerization mediated by the N-terminal helix (Guo *et al*., 2007; Kirsh *et al*., 2002). When HDAC4 accumulates in neuronal nuclei it forms larger punctate foci (Bolger *et al*., 2007; Lang *et al*., 2019; Sen *et al*., 2015), which have been variously described as speckles or aggregates. HDAC4 also co-aggregates with protein aggregates/inclusions that have been implicated in neurodegenerative diseases including ataxin-1 aggregates in cultured neurons (Bolger *et al*., 2007), and ubiquitinated intranuclear inclusions in the brains of individuals with Parkinson’s (Takahashi-Fujigasaki & Fujigasaki, 2006) and neuronal intranuclear inclusion diseases (Takahashi-Fujigasaki *et al*., 2006). HDAC4-containing aggregates in the cytoplasm have also been associated with disease; cytoplasmic HDAC4 is sequestered into mutant huntingtin aggregates in Huntington’s disease mouse models (Mielcarek *et al*., 2013a) and into cytoplasmic Lewy bodies in Parkinson’s disease (Takahashi-Fujigasaki & Fujigasaki, 2006). Mutations within the 14-3-3 binding site of HDAC4 have been identified in seven unrelated individuals with intellectual disability and developmental delay, and these mutations reduce the affinity of HDAC4 for 14-3-3 in cultured cells (Wakeling *et al*., 2021). Consequently, it is hypothesized that disrupted nucleocytoplasmic shuttling of HDAC4 underlies neuronal dysfunction in these individuals. Despite previously being referred to as aggregates, recent evidence indicates that HDAC4 foci are more accurately described as biomolecular condensates (termed condensates herein, Liu *et al*. (2024)), which are dynamic subnuclear compartments that regulate molecular interactions (Nam & Gwon, 2023). Disruption of these condensates, as observed in neuronal disease, is therefore likely to contribute to neuronal dysfunction. Despite numerous observations of HDAC4 accumulation and condensation in neuronal disease, the molecular mechanisms driving condensate formation and their contribution to disease onset and/or progression remain unexplored, highlighting a critical gap in understanding.

Human and *Drosophila* HDAC4 are highly conserved, sharing 35% amino acid sequence identity and 59% similarity across the entire protein (Fitzsimons *et al*., 2013). Importantly, this conservation includes key regulatory motifs such as the NLS, NES, MEF2 binding region, catalytic site, ankyrin repeat binding region and serine residues that mediate nuclear exit when phosphorylated (Tan *et al*., 2024). HDAC4 is highly expressed throughout the *Drosophila* brain (Fitzsimons *et al*., 2013; Tan *et al*., 2024), including in the mushroom body, which is a key integration center for sensory information that shares many architectural features with the vertebrate cerebellum (Li *et al*., 2020). The intrinsic neurons of the mushroom body, known as Kenyon cells, exhibit nuclear condensate formation and impaired long-term memory upon expression of either human or *Drosophila* HDAC4 (Fitzsimons *et al*., 2013; Main *et al*., 2021). Developmental overexpression of HDAC4 disrupts both mushroom body and eye formation (Main *et al*., 2021; Schwartz *et al*., 2016; Tan *et al*., 2024), and the severity is exacerbated by nuclear accumulation of HDAC4 (Main *et al*., 2021; Tan *et al*., 2024). The mushroom body defects are significantly curtailed by mutation of the MEF2 binding site within HDAC4, which correlates with a significant decrease in nuclear localization of HDAC4 (Main *et al*., 2021; Tan *et al*., 2024). In summary, expression of either human or *Drosophila* HDAC4 results in formation of condensates in neuronal nuclei (Fitzsimons *et al*., 2013; Main *et al*., 2021; Tan *et al*., 2024), with a positive correlation observed between increased nuclear localization, condensate number, and the severity of defects in neurodevelopment (Tan *et al*., 2024). These findings demonstrate that *Drosophila* provides an effective model for the mechanistic analysis of HDAC4 condensation *in vivo*.

The propensity of HDAC4 to form condensates appears to stem from its glutamine-rich N-terminus, as deletion of residues 1-179 of human HDAC4 prevent nuclear condensate formation (Kirsh *et al*., 2002). This region mediates homo-oligomerization of HDAC4 through its N-terminal α-helix; in solution, the N-terminus of human HDAC4 forms an α-helix that self-associates in a four-helix bundle (Guo *et al*. (2007), Fig. 1A). Tetramerization is dependent on the formation of a small hydrophobic core, consisting of Leu89, Ile90 and Phe93, which is supported by glutamine-dominated polar interaction networks. Substitution of Phe93 for a polar residue abolishes tetramerization in solution, while mutation of nearby His97 to phenylalanine extends the core and stabilizes tetramerization (Guo *et al*., 2007). Furthermore, it was recently demonstrated that in the presence of DNA and MEF2, the N-terminal region of HDAC4 preferentially forms a dimer, with each HDAC4 molecule bound to a MEF2 dimer that associates with its cognate DNA molecule (Dai *et al*., 2024). It is unclear, however, if these interactions translate into the 3D-conformation of full-length HDAC4 and thus influence its condensation *in vivo*.

**Figure 1.**
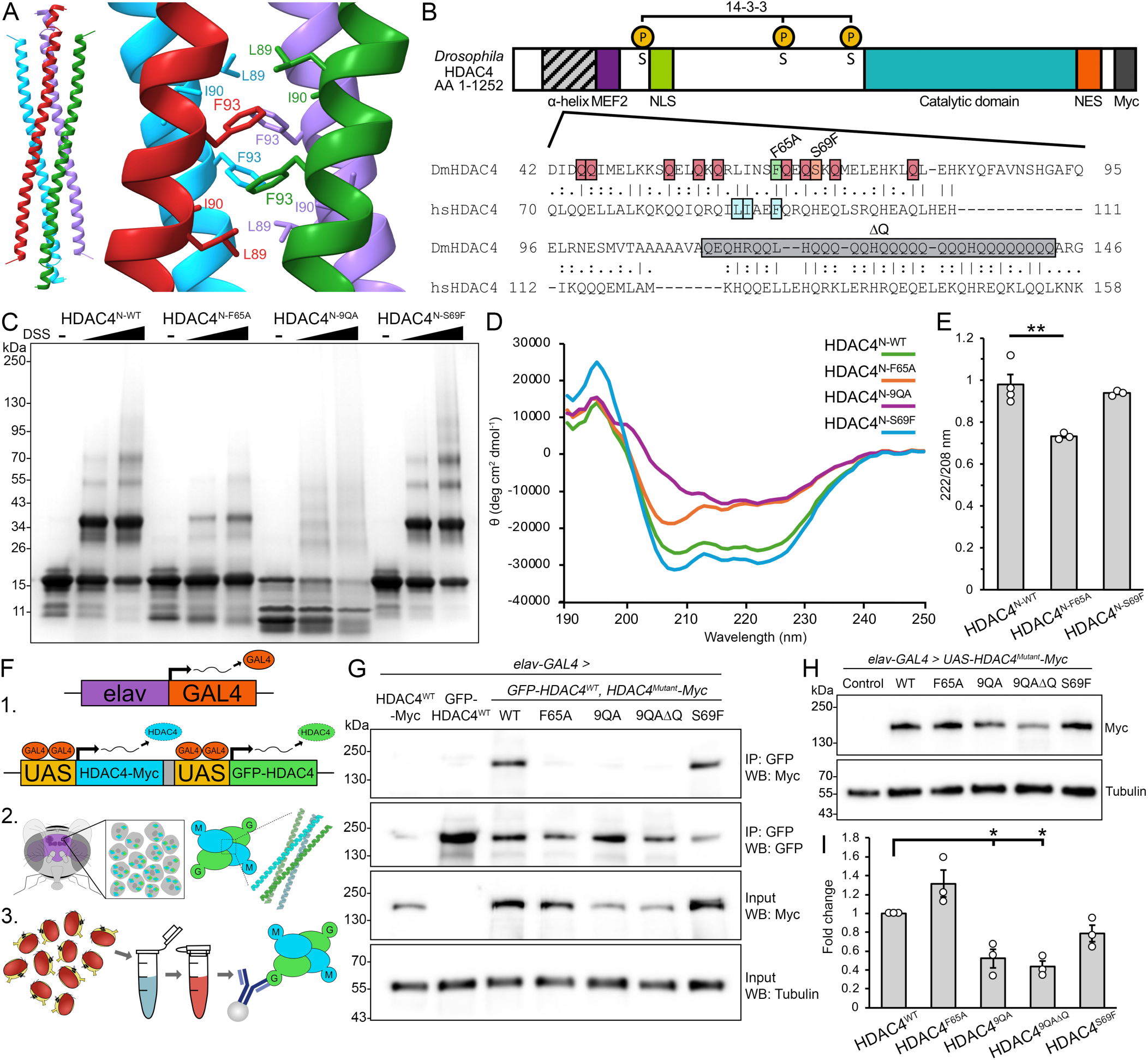
Mutation of the hydrophobic core or conserved glutamine residues alters oligomerization of DmHDAC4 in the adult brain. (A) Structure of the human HDAC4 (hsHDAC4) tetramer and hydrophobic core as determined by Guo et al. (2007) (PDB entry 2H8N). Left; the N-terminal region of hsHDAC4 (residues 62-153) forms an α-helix that assembles into a four-helix bundle, consisting of two pairs of antiparallel helices (pair one: green and red, pair two, blue and purple). Right; at the center of the tetramer is a hydrophobic core consisting of three nonpolar residues (L89, I90, F93), of which F93 is central. The orientations are presented with the same alignment and colour scheme as Guo et al. (2007). (B) Schematic diagram depicting the domain structure of Drosophila HDAC4 (top, DmHDAC4), including the N-terminal α-helix, MEF2 binding site (MEF2, purple) nuclear localization signal (NLS, green), 14-3-3 binding sites (P, phosphorylated serine (S) residues, yellow), catalytic domain (teal), nuclear export signal (NES, orange) and C-terminal Myc-tag (grey). Alignment of the α-helix region of hsHDAC4 and DmHDAC4 (bottom). Mutants used in this study: F65A, green box; 9QA, red boxes; 9QAΔQ, red boxes & grey ΔQ; S69F, orange box. The 3SA mutants substitute the three noted phosphorylated serines (S) for alanines. (C) SDS-PAGE of chemically crosslinked purified recombinant wild-type and mutant N-terminal HDAC4 (HDAC4^N^). In the absence of crosslinker (-, DMSO only) HDAC4^N^ appears as a ∼15 kDa monomer. Addition of 0.2 mM or 2.0 mM disuccinimidyl suberate (DSS) induces crosslinking, enabling visualization of dimer, trimer and tetramer species of HDAC4^N-WT^ and HDAC4^N-S69F^, while oligomerization is reduced for HDAC4^N-F65A^. HDAC4^N-9QA^ undergoes significant degradation. (D) Circular dichroism spectra of purified HDAC4^N^ variants. HDAC4^N-WT^ and HDAC4^N-S69F^ exhibit α-helical folding, with characteristic negative peaks at 208 and 222 nm. HDAC4^N-F65A^ shows a partial loss of α-helicity, while HDAC4^N-9QA^ exhibits a more pronounced reduction in α-helicity, indicating a shift toward a less ordered conformation. (E) 222/208 nm ratios from CD spectra of purified HDAC4^N^. HDAC4^N-WT^ and HDAC4^N-S69F^ maintain a ratio close to 1, consistent with oligomerization, whereas HDAC4^N-F65A^ shows a significant reduction, suggesting impaired oligomerization (n = 3 independent spectra). ANOVA, F_(2,7)_ = 13.77, p = 0.003747; post-hoc Tukey’s HSD, ** p < 0.01. Error bars indicate SEM. (F) Schematic of the co-immunoprecipitation workflow used in (G). 1. GFP- (green) and Myc-tagged (blue) HDAC4 are expressed in flies using the UAS/GAL4 system under the control of the pan-neuronal elav-GAL4 driver. 2. HDAC4 transgenes are expressed in the adult brain (purple), where they form condensates comprised of homo- and hetero-tetramers of GFP- and Myc-tagged HDAC4 in Kenyon cell nuclei. 3. Whole cell lysates are prepared from adult fly heads and subjected to immunoprecipitation. GFP-HDAC4 is immunoprecipitated from all samples and co-immunoprecipitated HDAC4-Myc (WT or mutant) is detected. (G) Mutation of the hydrophobic core or conserved glutamine residues alters oligomerization of HDAC4 in vivo. Genotypes were generated by crossing elav-GAL4 females to males carrying the indicated HDAC4 transgene. Flies were raised at 18 °C until eclosion when adults were transferred to 22 °C to increase transgene expression. Co-immunoprecipitation yield of HDAC4^F65A^ (lane 4), HDAC4^9QA^ (lane 5), and HDAC4^9QAΔQ^ (lane 6), is significantly reduced, and HDAC4^S69F^ (lane 7) increased, compared to HDAC4^WT^ (lane 3) upon immunoprecipitation of GFP-HDAC4. (H) Whole cell lysates generated from adult heads expressing HDAC4^WT^-Myc or HDAC4^Mutant^-Myc under the control of elav-GAL4 were subjected to SDS-PAGE and probed for Myc and tubulin. (I) Quantification of HDAC4 band intensity normalized to tubulin (n = 3 blots). HDAC4^9QA^ and HDAC4^9QAΔQ^ are significantly reduced compared to HDAC4^WT^. One sample t-test compared to HDAC4^WT^: HDAC4^F65A^ t_(2)_ = 2.201, p = 0.1586; HDAC4^9QA^ t_(2)_ = 4.8747, p = 0.0396; HDAC4^9QAΔQ^ t_(2)_ = 9.1283, p = 0.0118; HDAC4^S69F^ t_(2)_ = 2.3819, p = 0.1401.

Here we aimed to determine whether self-oligomerization of HDAC4 via the N-terminal α-helix is a requirement for condensate formation of the full-length protein, and to characterize the role of HDAC4 condensation in neuronal dysfunction in *Drosophila.* Informed by Guo *et al*. (2007) we expressed full-length HDAC4 mutants predicted to display altered ability to oligomerize and subsequently form condensates *in vivo*. Reduced condensation correlated with less severe overexpression-induced phenotypes in our *Drosophila* model, demonstrating that condensate formation contributes to pathogenesis. Oligomerization promotes nuclear import of HDAC4, but nuclear restricted HDAC4 oligomerization mutants were still impaired in their ability to form condensates. Preventing MEF2 binding further reduced condensation, suggesting that HDAC4 binding partners, including MEF2, stabilize condensate formation and exacerbate toxicity. Additionally, regions downstream of the MEF2 binding site also contribute to protein stabilization, condensate formation, and neurotoxicity. These findings emphasize the importance of understanding HDAC4 function, particularly regulated condensate formation, to better understand how its dysfunction contributes to neuronal disease.

## Results

### Characterization of HDAC4 oligomerization mutants

We first sought to examine whether the mechanism of oligomerization of *Drosophila* HDAC4 (DmHDAC4) is conserved with human HDAC4 (hsHDAC4). The sequence of the N-terminal α-helix of hsHDAC4 is highly conserved with DmHDAC4 (Fig. 1B). Notably, the three non-polar residues that form the hydrophobic core (Leu61, Ile62 and Phe65 in *Drosophila*) are all strictly conserved, as are nine of the glutamine residues that stabilize intra- and interhelical interactions (Guo *et al*., 2007). As tetramerization of hsHDAC4 is destabilized upon substitution of Phe93 within the hydrophobic core (Guo *et al*., 2007), we generated the homologous mutation within DmHDAC4, Phe65Ala (HDAC4^F65A^). Either side of this core are the glutamine residues that form intra- and inter-helical polar interaction networks. To investigate the importance of these networks for tetramer formation, the nine conserved glutamine residues were substituted with alanines (HDAC4^9QA^). While the N-termini of both hsHDAC4 and DmHDAC4 are glutamine-rich, DmHDAC4 has a significant polyglutamine stretch C-terminal to the nine conserved glutamines. Given the role that polyglutamine regions have in protein aggregation (Michalik & Van Broeckhoven, 2003), as well as the roles of glutamines in the potentially stabilizing interaction networks of the HDAC4 tetramer (Guo *et al*., 2007), this 32 amino acid stretch (residues Gln112-Gln143) containing 25 glutamine residues was deleted alongside the 9QA substitutions (HDAC4^9QAΔQ^) to determine whether it confers additional stability to DmHDAC4 oligomers. Conversely, since hsHDAC4 tetramers are stabilized by extension of the hydrophobic core via substitution of His97 to Phe (Guo *et al*., 2007), we also generated the corresponding Ser69Phe (HDAC4^S69F^) mutation.

#### In vitro oligomerization

To characterize the capacity of these mutants to oligomerize, *in vitro* chemical crosslinking experiments were performed using purified recombinant N-terminal fragments spanning Pro37 to Gln143 of DmHDAC4, which corresponds to the region of hsHDAC4 examined by Guo *et al*. (2007). To avoid confusion with full-length HDAC4 constructs, these are denoted as HDAC4^N-mutant^. HDAC4^N-WT^ formed oligomers corresponding to homodimer, trimer, and tetramers as well as higher order species (Fig. 1C). HDAC4^N-F65A^ showed a significant reduction across all oligomeric species, whereas HDAC4^N-S69F^ exhibited a slight increase in trimers and tetramers, with higher order species appearing more prominent. In contrast, significant degradation of HDAC4^N-9QA^ occurred during affinity and size-exclusion purification, however the presence of higher molecular weight smears on SDS-PAGE suggest that oligomerization still occurred, though the resulting species did not correspond to those observed for HDAC4^N-WT^. Changes in oligomerization were also reflected by changes in retention volume between the mutants and wild-type during size-exclusion chromatography purification (SFig. 1), as observed with hsHDAC4 (Dai *et al*., 2024).

Circular dichroism analysis revealed that HDAC4^N-WT^ primarily adopted an α-helical secondary conformation in solution, similarly to hsHDAC4 (Guo *et al*., 2007), as did HDAC4^N-F65A^ and HDAC4^N-S69F^ (Fig. 1D). However, this helical conformation was lost for HDAC4^N-9QA^, suggesting it exists as a less structured or partially unfolded helix. Comparison of the 222/208 nm ratio for structures that form an α-helix provides insight into helical packing and oligomerization of proteins, where ratios close to or above 1 suggest coiled-coil or tightly packed helices, and lower values indicate more isolated helices or structural destabilization (Crooks *et al*., 2011). Both HDAC4^N-WT^ and HDAC4^N-S69F^ had ratios close to 1 (Fig. 1E), supporting that they exist as oligomeric helices, while the ratio for HDAC4 ^N-F65A^ was reduced, suggestive of forming more isolated helices. These data indicate that F65A stabilises and S69F destabilises HDAC4 oligomers, while glutamine residues are required for the correct α-helical folding of HDAC4.

#### In vivo oligomerization

To characterize the ability of each of the full-length HDAC4 mutants to oligomerize *in vivo*, transgenic flies were generated carrying UAS-HDAC4^WT^ or each of the mutants (hereafter collectively referred to as HDAC4^Mutant^) for UAS/GAL4 regulated expression in the brain (Brand & Perrimon, 1993). To determine whether the HDAC4 mutants retained the capacity to oligomerize, co-immunoprecipitation was performed on whole head lysates from flies in which UAS-GFP-HDAC4^WT^ (Main *et al*., 2021) was pan-neuronally co-expressed with either UAS-HDAC4^WT^-Myc or UAS-HDAC4^Mutant^-Myc via the *elav-GAL4* driver (Hawley *et al*., 2023; Robinow & White, 1988), (Fig. 1F). Immunoprecipitation of GFP-HDAC4^WT^ resulted in co-immunoprecipitation of HDAC4^WT^-Myc (Fig. 1G, lane 3), confirming oligomerization of wild-type DmHDAC4. HDAC4^S69F^ (Fig. 1G, lane 7) co-immunoprecipitated similarly to HDAC4^WT^, while HDAC4^F65A^, HDAC4^9QA^ and HDAC4^9QAΔQ^ were all significantly reduced in co-immunoprecipitation yield (Fig. 1G, lanes 4-6), revealing a severely compromised ability to oligomerize. It was noted however via quantitative analysis of whole head lysates that the amount of HDAC4^9QA^ and HDAC4^9QAΔQ^ was reduced to approximately half that of HDAC4^WT^ (Fig. 1H-I, p < 0.001), indicating that as well as mediating oligomerization and stabilizing the N-terminal helix of HDAC4, the glutamine residues may be important in full-length HDAC4 folding, where their loss destabilises protein levels. Together these data demonstrate that both the hydrophobic core and glutamine-mediated polar interaction networks and/or α-helical folding are essential for oligomerization of the N-terminus of HDAC4 *in vitro*, and oligomerization of full-length HDAC4 *in vivo*. For simplicity, the multimerization of HDAC4 is herein referred to as oligomerization, as the exact structural form *in vivo* has not been determined (i.e. it is unclear whether multimers represent dimers, trimers, tetramers, or higher order species *in vivo*).

### Oligomerization of HDAC4 promotes nuclear condensate formation in mushroom body neuronal nuclei

We next examined whether the impaired ability of HDAC4 to oligomerize led to reduced nuclear condensate formation *in vivo*. *UAS-HDAC4-Myc* expression was induced in the mushroom body using the *OK107-GAL4* driver (Connolly *et al*., 1996; Hawley *et al*., 2023). To ensure that transgene expression did not impact the development of these cells, expression was induced in the adult brain via the TARGET system in which GAL4 is repressed during development at 18°C by a temperature-sensitive mutant of GAL80 (GAL80ts). Once adult flies emerged, they were transferred to 30°C, at which temperature GAL80ts repression of GAL4 is relieved and HDAC4 transgene expression is permitted (McGuire *et al*. (2004), Fig. 2A). MEF2 was used as a counterstain to enhance the visibility of HDAC4 condensates as it is sequestered in HDAC4 nuclear foci (Chan *et al*., 2003; Fitzsimons *et al*., 2013; Miska *et al*., 1999; Tan *et al*., 2024). HDAC4^WT^ was robustly expressed in adult mushroom body neurons (Fig. 2B,C), and formed nuclear condensates that colocalized with MEF2 (Fig. 2E), as observed previously (Fitzsimons *et al*., 2013; Tan *et al*., 2024). Expression of HDAC4^F65A^, HDAC4^9QA^ and HDAC4^9QAΔQ^ resulted in significantly fewer and visibly smaller condensates than HDAC4^WT^, whereas HDAC4^S69F^ expression doubled the number of condensates. (Fig. 2D). These data collectively demonstrate a positive correlation between the HDAC4 N-terminal α-helix’s oligomerization capacity and condensate formation *in vivo*. Although condensate formation was significantly decreased by HDAC4^F65A^, HDAC4^9QA^ and HDAC4^9QAΔQ^, it was not completely abolished. We hypothesize this is due to the incomplete loss of oligomerization in each mutant (Fig. 1C). Additionally, the presence of endogenous HDAC4 likely contributes to oligomerization and subsequent condensate formation; we previously confirmed that endogenous HDAC4 forms condensates with transgene HDAC4^WT^ (Tan *et al*., 2024).

**Figure 2.**
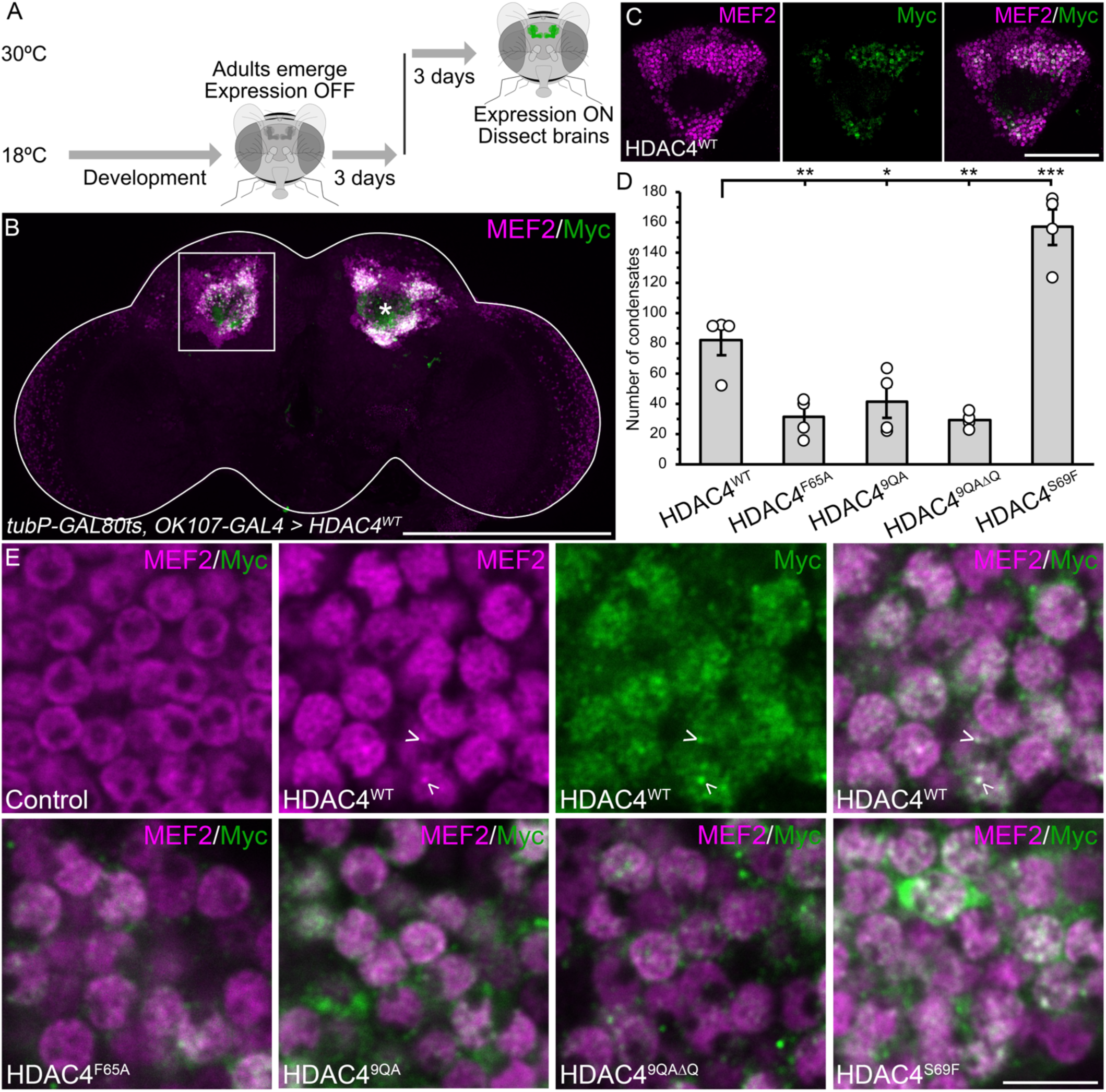
HDAC4 oligomerization mutants are altered in their ability to form nuclear condensates. All genotypes were generated by crossing *tubP-GAL80ts; OK107-GAL4* females to males carrying the indicated *UAS-HDAC4-Myc* transgene and to the *w(CS10)* control, and transgene expression was induced in adulthood with GAL80ts. (A) Schematic of the protocol for induction of gene expression in the adult mushroom body. Flies were raised at 18 °C, at which temperature GAL80ts represses GAL4, until after eclosion when adult flies were shifted to 30 °C. At this temperature GAL80ts is inactivated, enabling GAL4-induced gene expression. Brains were dissected after 72 hours at 30 °C. (B-C, E) Whole mount brains were subjected to immunohistochemistry with anti-Myc (green) and anti-MEF2 (magenta). (B) Maximum projection through the posterior of the whole brain demonstrating *OK107-GAL4*-mediated expression of HDAC4^WT^-Myc, which colocalizes with endogenous MEF2 in mushroom body neuronal nuclei. Asterisk labels the calyx and the boxed region is depicted in (C). Scale bar = 200 µm. (C) Single optical section (0.5 µm) through the calyx and Kenyon cell layer of the posterior of the brain showing HDAC4^WT^ expression and colocalization with MEF2. Scale bar = 50 µm. (D) Quantification of nuclear condensation of HDAC4. The number of condensates (colocalizing puncta of HDAC4 and MEF2) per optical section (as shown in (C)) were counted and averaged for 10 sections through the posterior of the brain at 1 µm increments (n = 4 brains per genotype). ANOVA, F_(4,15)_ = 36.88, p < 0.0001; post-hoc Tukey’s HSD, * p < 0.05, ** p < 0.01, *** p < 0.001. Error bars indicate SEM. (E) Representative images of HDAC4 condensates in mushroom body neurons. Arrowheads point to nuclear condensates of HDAC4. Single optical sections (0.5 µm) are shown. Scale bar = 5 µm.

### HDAC4 forms dynamic condensates in vivo

Our data clearly demonstrate that N-terminal oligomerization is required for nuclear condensation of HDAC4. However, it remained uncertain whether downstream regions also contribute to this process. To that end, we generated HA-tagged full-length HDAC4 alongside a truncated N-terminal construct of HDAC4 comprising residues Met1 to Leu285 (HDAC4^M1-L285^), which includes the MEF2 binding site to facilitate nuclear import (Fig. 3A). One of the serine residues important in 14-3-3 binding is incorporated in this region (Ser 239), therefore we mutated this to alanine to prevent nuclear exit. When expression was induced in the adult mushroom body under the same conditions as in Fig. 1, HDAC4^WT^ was observed in both cytoplasmic haloes and in the nucleus, where it formed condensates (Fig. 3B). In contrast, HDAC4^M1-L285^ localised exclusively to the nucleus, where it appeared more uniform and granular than HDAC4^WT^, and did not form condensates containing MEF2. Constitutive expression throughout development (i.e. in the absence of GAL80ts) resulted in larger HDAC4^WT^ condensates, but still no condensate formation was observed for HDAC4^M1-L285^ (Fig. 3C). Similarly, no condensates formed when expressed using the *elav-GAL4* driver (Fig. 3D). We confirmed that this was not a result of differing protein levels as although HDAC4^WT^ total protein was four times higher than HDAC4^M1-L285^ in whole head lysates (Fig. 3E-F), both had a similar concentration localised to nuclear fractions (Fig. 3G-H). Together these data reveal that while oligomerization of the N-terminal region of HDAC4 is necessary for condensation, C-terminal regions are also required. These data also validate that condensate formation is not due to the presence of the Myc tag, and further we have also observed condensates with FLAG- (Fitzsimons *et al*., 2013) and GFP- (Main *et al*., 2021) tagged HDAC4 constructs.

**Figure 3.**
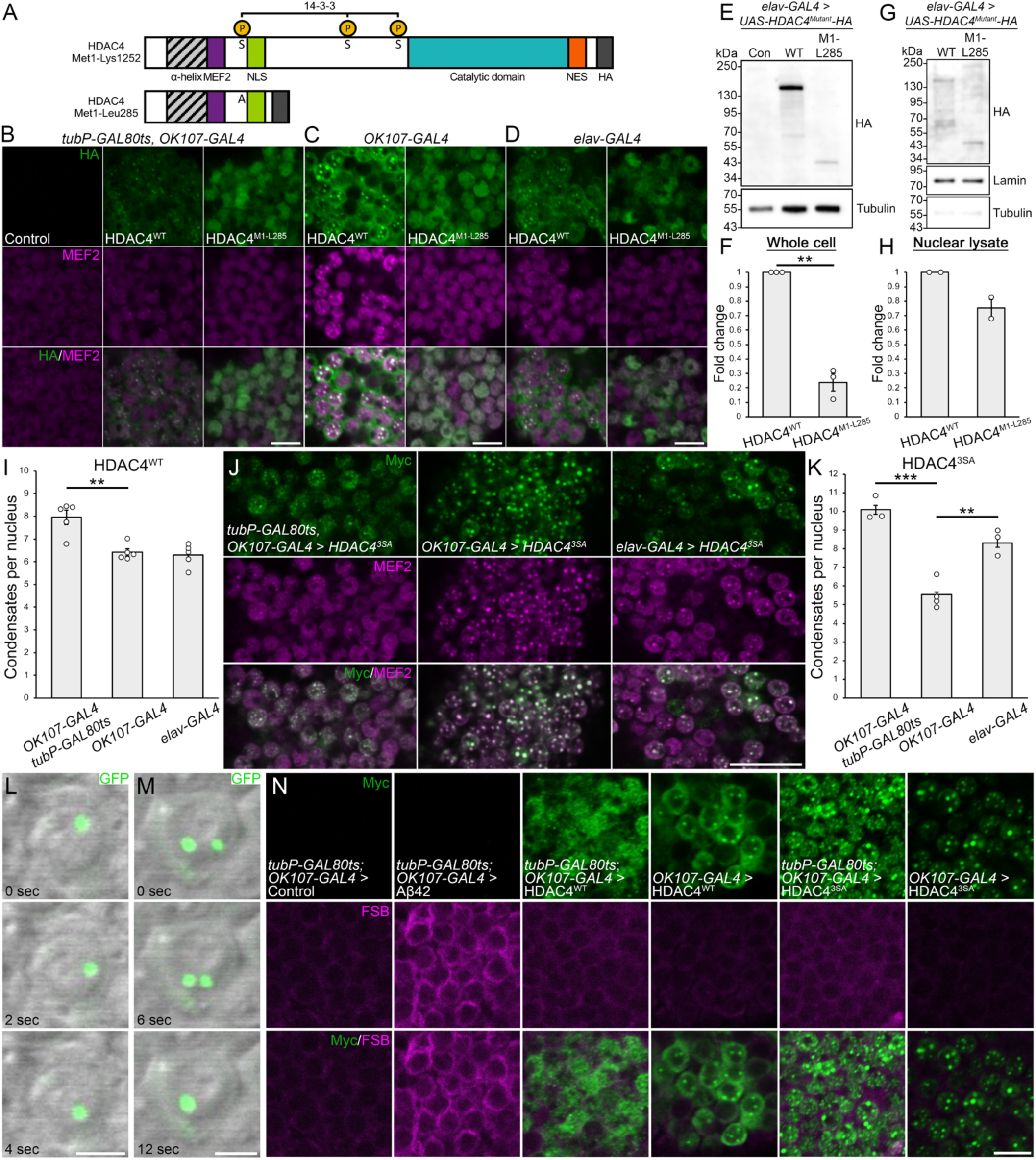
HDAC4 condensate formation requires regions downstream of the N-terminus and is dynamic. (A) Schematic diagram depicting the domain structure of full-length DmHDAC4 (top) and the truncated HDAC4^M1-L285^ (bottom). Both share the N-terminal α-helix, MEF2 binding site (MEF2, purple) nuclear localization signal (NLS, green) and a C-terminal HA-tag (grey), but only full-length HDAC4 contains intact 14-3-3 binding sites (P, phosphorylated serine (S) residues), catalytic domain (teal) and nuclear export signal (NES, orange). The first serine (S239) is mutated to alanine in HDAC4^M1-L285^. (B-D) Whole mount brains were subjected to immunohistochemistry with anti-Myc (green) and anti-MEF2 (magenta). Representative single optical sections (0.5 µm) through the Kenyon cell layer are shown. Under all conditions HDAC4^WT^ formed visible condensates, but these were not observed for HDAC4^M1-L285^, which instead appeared granular. Scale bar = 5 µm. (B) Genotypes were generated by crossing *tubP-GAL80ts; OK107-GAL4* females to males carrying either *UAS-HDAC4^WT^-HA* or *UAS-HDAC4^M1-L285^-HA*. Flies were raised at 18 °C throughout development and transgene expression was induced by incubation of flies at 30 °C for 72 hours. (C-D) Genotypes were generated by crossing *OK107-GAL4* (C) or *elav-GAL4* (D) females to males carrying either *UAS-HDAC4^WT^-HA* or *UAS-HDAC4^M1-L285^-HA*. Flies were raised at 25 °C throughout development and adulthood. (E) Whole cell lysates generated from adult heads expressing HDAC4^WT^ or HDAC4^M1-L285^ under the control of *elav-GAL4* were subjected to western blotting and probed for Myc and tubulin. (F) Quantification of HDAC4 band intensity (from E) normalized to tubulin (n = 3). HDAC4^M1-L285^ is significantly reduced compared to HDAC4^WT^. One sample t-test, t_(2)_ = 12.633, p = 0.0062. Error bars indicate SEM. (G) Subcellular fractionation and western blotting was performed on fly heads. Membranes were probed with anti-Myc, as well as anti-lamin and anti-α-tubulin to assess fractionation efficacy. (H) Quantification of nuclear HDAC4 (from G), normalized to lamin (n = 2). There is no significant difference in nuclear abundance between HDAC4^WT^ and HDAC4^M1-L285^. One sample t-test, t_(1)_ = 3.348, p = 0.185. Error bars indicate SEM. (I) Quantification of HDAC4^WT^ nuclear condensation. The number of condensates (colocalizing puncta of HDAC4 and MEF2) per nucleus were counted and averaged for n ≥ 28 nuclei per section for n = 5 brains per genotype. ANOVA, F_(2,12)_ = 14.67, p = 0.000599; post-hoc Tukey’s HSD, ** p < 0.01. Error bars indicate SEM. (J) Whole mount brains were subjected to immunohistochemistry with anti-Myc (green) and anti-MEF2 (magenta). Representative single optical sections (0.5 µm) through the Kenyon cell layer are shown. Genotypes were generated by crossing *tubP-GAL80ts; OK107-GAL4, OK107-GAL4*, or *elav-GAL4* females to males carrying UAS-HDAC4^3SA^-Myc. All flies were raised at 18 °C throughout development and transgene expression was induced/increased by incubation of flies at 30 °C for 72 hours. HDAC4^3SA^ condensate formation varied depending on expression conditions. Scale bar = 10 µm. (K) Quantification of nuclear condensation of HDAC4^3SA^. The number of condensates (colocalizing puncta of HDAC4 and MEF2) per nucleus were counted and averaged for n ≥ 27 nuclei per section for n ≥ 3 brains per genotype. ANOVA, F_(2,7)_ = 36.07, p = 0.000206; post-hoc Tukey’s HSD, ** p < 0.01, *** p < 0.001. Error bars indicate SEM. (L-M) live imaging performed on whole mount brains. Genotypes were generated by crossing *OK107-GAL4* females to *UAS-HDAC4^WT^-GFP* males, and raised at 18 °C throughout development and adulthood. Differential interference contrast (DIC) microscopy was used to visualise nuclei (gray) and intrinsic GFP fluorescence was detected (green). (L) HDAC4^WT^-GFP condensates are mobile within nuclei. (M) Coalescence of two HDAC4^WT^-GFP condensates. Scale bar = 2 µm. (N) Whole mount brains were subjected to immunohistochemistry with anti-Myc (green) and then stained with 1-Fluoro-2,5-bis(3-carboxy-4-hydroxystyryl)benzene (FSB, magenta). Representative single optical sections (0.5 µm) through the Kenyon cell layer are shown. Genotypes were generated as described in (B, C). FSB successfully detected Aβ42, but did not label HDAC4 condensates. Scale bar = 5 µm.

We also observed that constitutive expression of HDAC4^WT^ led to an increase in condensate size (compare Fig. 3B and C), which was accompanied by a noticeable decrease in condensate number (Fig. 3I), suggesting that prolonged expression drives coalescence. However, HDAC4^WT^ retains the ability to shuttle between the nucleus and cytoplasm, and it is unclear whether this is modulated by level of expression. To explore dose-dependent HDAC4 condensation further and eliminate any potential changes in subcellular distribution, we examined the condensates formed by expression of the HDAC4^3SA^ mutant. This mutant carries three serine to alanine substitutions (Ser239Ala, Ser573Ala and Ser748Ala) that prevent 14-3-3 binding and subsequent nuclear export, thus confining it to the nucleus (Tan *et al*., 2024; Wang *et al*., 2000; Wang & Yang, 2001). All flies were raised at 18°C until adulthood and then transferred to 30°C to induce transgene expression. *OK107-GAL4* drives a higher level of expression in mushroom body neurons than *elav-GAL4* (Hawley *et al*., 2023), and under the control of *elav-GAL4*, total HDAC4^3SA^ levels were decreased compared to *OK107-GAL4* (SFig. 2A). Reducing the level of expression in Kenyon cells decreased HDAC4^3SA^ condensate size (Fig. 3J), but increased the number of condensates per nucleus (Fig. 3K). The total protein level of HDAC4^3SA^ did not differ when expressed with *OK107-GAL4* in the presence or absence of GAL80ts (SFig. 2B), however condensate size increased and the number of condensates per nucleus decreased in the absence of GAL80ts. We hypothesized these changes resulted from condensate coalescence, therefore live-imaging of HDAC4^WT^-GFP expressed under the control of *OK107-GAL4* was performed. HDAC4 condensates were highly mobile (Fig. 3L) and underwent fusion events (Fig. 3M). Together these data demonstrate that HDAC4 condensation is a dynamic dose-dependent process, wherein condensate coalescence occurs following increasing protein levels, resulting in decreased condensate number but increased size.

Finally, we examined colocalization of HDAC4 condensates with the Congo red derivative FSB (1-Fluoro-2,5-bis(3-carboxy-4-hydroxystyryl)benzene) and Thioflavin-T (THT), markers of β-sheet amyloid aggregation (Juhl *et al*., 2019). FSB and THT successfully detected the 42 amino acid Aβ fragment of human Amyloid Precursor Protein in Kenyon cells when expressed under the control of *OK107-GAL4* (Fig. 3N, SFig. 3). However, neither FSB nor THT colocalized with HDAC4^WT^ condensates, regardless of the presence or absence of GAL80ts, nor with nuclear HDAC4^3SA^. This suggests that these protein accumulations are not dominated by amyloid cross β-sheet structures.

These findings highlight that HDAC4 condensate formation depends on both N-terminal oligomerization and C-terminal protein domains. This condensation occurs in a dynamic manner, influenced by expression level and timing, and lacks amyloid-like characteristics, suggesting that HDAC4 oligomerization and accumulation may have significant biological relevance. We continue to refer to HDAC4 inclusions as biomolecular condensates, given their dependence on N-terminal self-polymerization for their formation and their characteristics indicating that they likely form via liquid-liquid phase separation, including concentration-dependent foci formation and coalescence.

### Condensation of HDAC4 correlates with the severity of HDAC4 overexpression-induced neurodevelopmental defects

Building on these findings, we next examined whether the capacity of each mutant to form condensates was linked to the severity of neurodevelopmental defects induced by HDAC4 in the mushroom body. The *Drosophila* mushroom body is a bilateral structure composed of bundled axons and thus is ideal for monitoring gross changes in axon morphogenesis. These axons project from Kenyon cell bodies in the posterior of the brain in a bundle termed the pedunculus before bifurcating based on their subtype (α/β, α’/β’, and γ) to form characteristic lobes (Fig. 4A), (Lee *et al*., 1999). The cell adhesion molecule fasciclin II (FasII) is abundantly expressed in the α/β and γ lobes, allowing for clear visualization of these lobes when immunostained with anti-FasII (Freymuth & Fitzsimons, 2017). Expression of HDAC4^WT^-Myc with the pan-neuronal *elav-GAL4* driver results in various mushroom body defects including premature lobe termination (missing lobe), lobe thinning and lobe fusion (Fig. 4A, Table 1), as observed previously (Main *et al*., 2021; Tan *et al*., 2024). These defects occurred at a similar rate upon expression of HDAC4^WT^ with an HA-epitope tag, but no defects were observed for HDAC4^M1-L285^ with the same tag (Fig. 4B), further illustrating the relationship between these defects and condensation of HDAC4.

**Figure 4.**
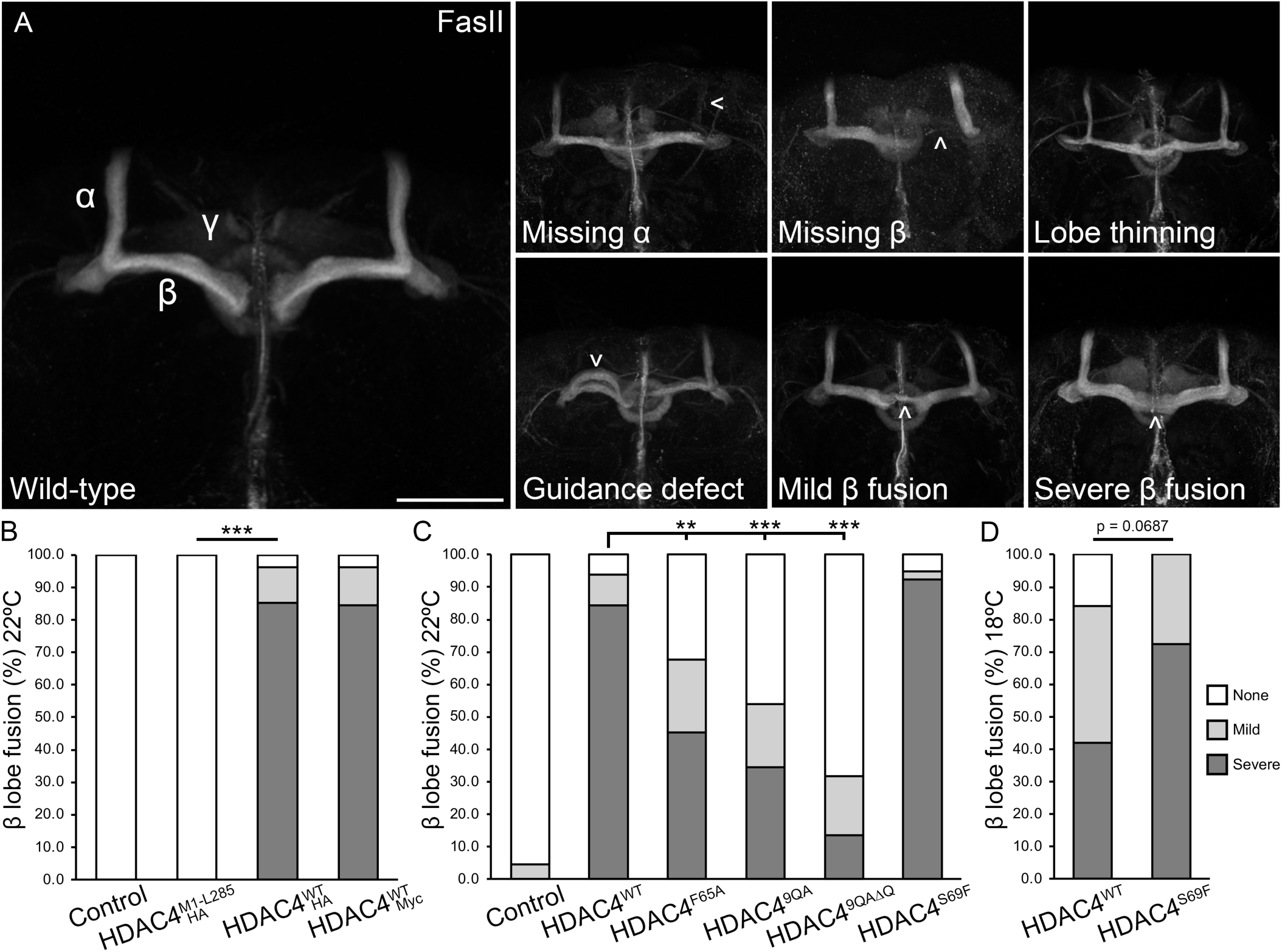
HDAC4 oligomerization correlates with severity of defects in mushroom body development. (A) Whole mount brains were subjected to immunohistochemistry with anti-Fasciclin II (FasII) to label alpha (α), beta (β), and gamma (γ) lobes of the adult mushroom body. Maximum projections were generated from stacks acquired at 1 µm increments. Defects (arrows) resulting from HDAC4-overexpression include lobe absence, α and β lobe thinning, guidance defects, mild β lobe fusion, and severe β lobe fusion. Scale bar 50 = µm. (B-D) Quantification of β lobe fusion resulting from HDAC4 overexpression. All genotypes were generated by crossing *elav-GAL4* females to males carrying the indicated *UAS-HDAC4* transgene or the *w(CS10)* control. Mild fusion is counted when less than half of the β lobe is fused, and severe when fusion is greater than half. Significance was determined using the Fisher’s exact test, ** p < 0.01, *** p < 0.001. (B) HDAC4^WT^-Myc and HDAC4^WT^-HA both induce β lobe fusion to a similar degree, while no fusion was observed for HDAC4^M1-L285^-HA. (C) The proportion of brains displaying β lobe fusion was significantly reduced for HDAC4^F65A^, HDAC4^9QA^ and HDAC4^9QAΔQ^ mutants. (D) Decreasing expression level exacerbates the difference in severity between HDAC4^WT^ and HDAC4^S69F^ but this was not to statistical significance.

**Table 1.** Full-length HDAC4 overexpression induces defects in mushroom body development. All genotypes were generated by crossing *elav-GAL4* females to males carrying each *UAS-HDAC4* transgene and to the *w(CS10)* control. Flies were raised at 22 °C. The percentage of brains displaying each phenotype was calculated from the total number of brains analysed for each genotype (n). Statistical analysis was performed with the Fisher’s Exact Test. Overexpression of HDAC4^WT^-Myc and HDAC4^WT^-HA each induced severe β lobe fusion compared to control (p < 0.0001), but HDAC4^M1-L285^ did not (p = 1).

| 22 °C |  |  |  |  |
| --- | --- | --- | --- | --- |
| (% of brains) | Control | HDAC4<sup>M1-L285</sup>-HA | HDAC4<sup>WT</sup>-HA | HDAC4<sup>WT</sup>-Myc |
| $\alpha$ lobe thinning | 4 | 0 | 59 | 35 |
| $\alpha$ lobe missing | 0 | 0 | 0 | 4 |
| $\beta$ lobe thinning | 0 | 0 | 33 | 0 |
| $\beta$ lobe missing | 0 | 0 | 0 | 4 |
| $\beta$ lobe fusion | 0 | 0 | 97 | 96 |
| Mild | 0 | 0 | 12 | 11 |
| Severe | 0 | 0 | 85 | 85 |
| Guidance defect | 0 | 0 | 4 | 0 |
| No defects | 100 | 100 | 0 | 4 |
| n | 23 | 22 | 27 | 26 |

The most prominent defect was β lobe fusion, which results from a failure of the β lobe axons to correctly terminate, resulting in erroneous crossing of the midline. β lobe fusion occurred in the majority of HDAC4^WT^ brains, which was significantly reduced by HDAC4^F65A^, HDAC4^9QA^ and HDAC4^9QAΔQ^ (Fig. 4C, Table 2). Subclassification of β lobe fusion to either mild (less than half of the lobe fused) or severe (more than half of the lobe fused) revealed that the mutants also reduced the severity of the phenotype (Table 2). Given the almost complete penetrance of β lobe fusion resulting from expression of HDAC4^WT^, it was unsurprising that there was no significant increase upon expression of HDAC4^S69F^. The severity of β lobe fusion correlates with the level of expression (Tan *et al*., 2024). Therefore, we reduced expression by lowering the temperature, which decreases GAL4 activity (Duffy, 2002), to uncover any differences between HDAC4^WT^ and HDAC4^S69F^. This exacerbated the difference in β lobe fusion between HDAC4^WT^ (84% total, 42% severe) and HDAC4^S69F^ (100% total, 72% severe), respectively, however this did not quite reach statistical significance (p = 0.0687) (Table 2, Fig. 4D). Nevertheless, together these data together demonstrate a correlation between nuclear condensation of HDAC4 and the severity of β lobe fusion in the mushroom body. Since neither HDAC4^9QA^ or HDAC4^F65A^ completely eliminated condensation or mushroom body defects, we combined HDAC4^9QA^ and HDAC4^F65A^ mutations as we hypothesized this may further destabilize oligomerization and subsequent HDAC4 condensation, however HDAC4^9QA-F65A^ did not further reduce β lobe fusion (STable 1) despite a small but insignificant reduction in condensate number (SFig. 4).

**Table 2.**
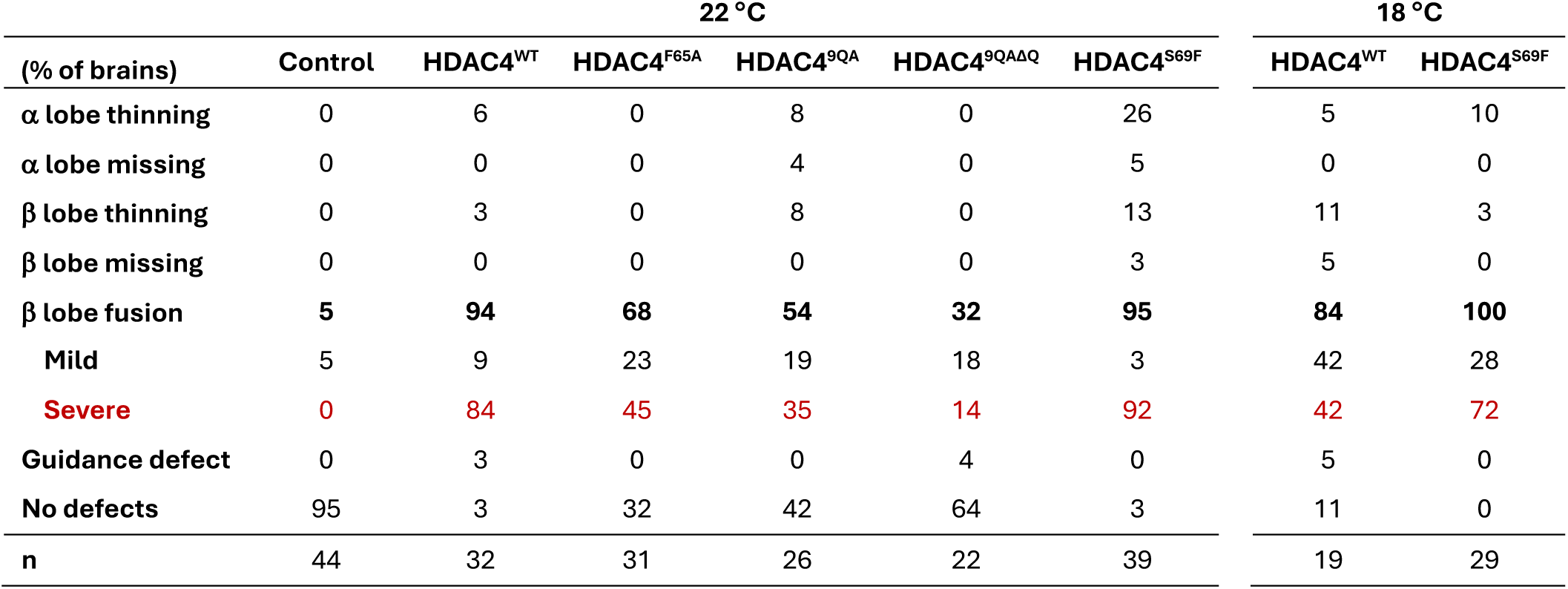
HDAC4 oligomerization mutants reduce HDAC4-overexpression induced defects in mushroom body development. All genotypes were generated by crossing *elav-GAL4* females to males carrying each *UAS-HDAC4-Myc* transgene and to the *w(CS10)* control. The percentage of brains displaying each phenotype was calculated from the total number of brains analysed for each genotype (n). Statistical analysis was performed with the Fisher’s Exact Test. Overexpression of HDAC4^WT^ resulted in a significant number of brains with β lobe fusion compared to control (p < 0.0001). Expression of HDAC4^F65A^ significantly reduced β lobe fusion compared to HDAC4^WT^ (p = 0.0017), as did expression of HDAC4^9QA^ (p < 0.0001) and HDAC4^9QAΔQ^ (p < 0.0001). HDAC4^S69F^ did not significantly increase the severity of all (p = 1) and severe (p = 0.3377) β lobe fusion compared to HDAC4^WT^. Percentages may not sum to 100 due to rounding.

| 22 °C |  |  |  |  |  |  | 18 °C |  |
| --- | --- | --- | --- | --- | --- | --- | --- | --- |
| (% of brains) | Control | HDAC4<sup>WT</sup> | HDAC4<sup>F65A</sup> | HDAC4<sup>9QA</sup> | HDAC4<sup>9QAΔQ</sup> | HDAC4<sup>S69F</sup> | HDAC4<sup>WT</sup> | HDAC4<sup>S69F</sup> |
| $\alpha$ lobe thinning | 0 | 6 | 0 | 8 | 0 | 26 | 5 | 10 |
| $\alpha$ lobe missing | 0 | 0 | 0 | 4 | 0 | 5 | 0 | 0 |
| $\beta$ lobe thinning | 0 | 3 | 0 | 8 | 0 | 13 | 11 | 3 |
| $\beta$ lobe missing | 0 | 0 | 0 | 0 | 0 | 3 | 5 | 0 |
| $\beta$ lobe fusion | 5 | 94 | 68 | 54 | 32 | 95 | 84 | 100 |
| Mild | 5 | 9 | 23 | 19 | 18 | 3 | 42 | 28 |
| Severe | 0 | 84 | 45 | 35 | 14 | 92 | 42 | 72 |
| Guidance defect | 0 | 3 | 0 | 0 | 4 | 0 | 5 | 0 |
| No defects | 95 | 3 | 32 | 42 | 64 | 3 | 11 | 0 |
| n | 44 | 32 | 31 | 26 | 22 | 39 | 19 | 29 |

Overexpression of HDAC4 is also associated with impaired eye development in *Drosophila* (Main *et al*., 2021; Schwartz *et al*., 2016; Tan *et al*., 2024). Development of the *Drosophila* eye initiates in early larval development and this process has been well characterized (Cagan, 2009; Cagan & Ready, 1989; Kumar, 2012). Overexpression of HDAC4^WT^ governed by the *glass multimer reporter* driver (*GMR-GAL4*) in post-mitotic photoreceptors (Freeman, 1996) disrupts development of the eye (Main *et al*., 2021; Schwartz *et al*., 2016; Tan *et al*., 2024), resulting in significant fusion, dimpling, and disorganization of ommatidia, as well as the loss of pigmentation, and the absence of bristles (Fig. 5A,B). The same disruption to development is observed for HDAC4^WT^-HA but not for HDAC4^M1-L285^-HA (SFig. 5). Expression of HDAC4^9QA^ and HDAC4^9QAΔQ^ resulted in significantly milder defects, as assessed via a semi-quantitative scoring system (Fig. 5C). Strikingly, expression of HDAC4^F65A^ had no effect on eye development, with ommatidia appearing wild type with normal pigmentation. In contrast, expression of HDAC4^S69F^ exacerbated the defects with almost complete fusion of all ommatidia and formation of cavities (Fig. 5A,B). To directly correlate changes in severity of the eye phenotype with condensation of HDAC4, immunohistochemistry was performed on the third instar larval eye disc (Fig. 5D). Nuclear condensation of HDAC4^WT^ occurred in the basal cells of the disc (Fig. 5E,F) as we observed previously (Tan *et al*., 2024). In contrast, the HDAC4^F65A^, HDAC4^9QA^ and HDAC4^9QAΔQ^ mutants that reduced the severity of the eye phenotype formed much fewer and smaller nuclear condensates, appearing predominantly in cytoplasmic haloes, while HDAC4^S69F^ formed more condensates (Fig. 5F). Together these data strengthen the evidence that oligomerization of HDAC4 via the N-terminal α-helix is the primary mechanism through which HDAC4 forms condensates, and that condensation of HDAC4 in the nucleus correlates with the severity of neurodevelopmental defects in both the *Drosophila* brain and eye.

**Figure 5.**
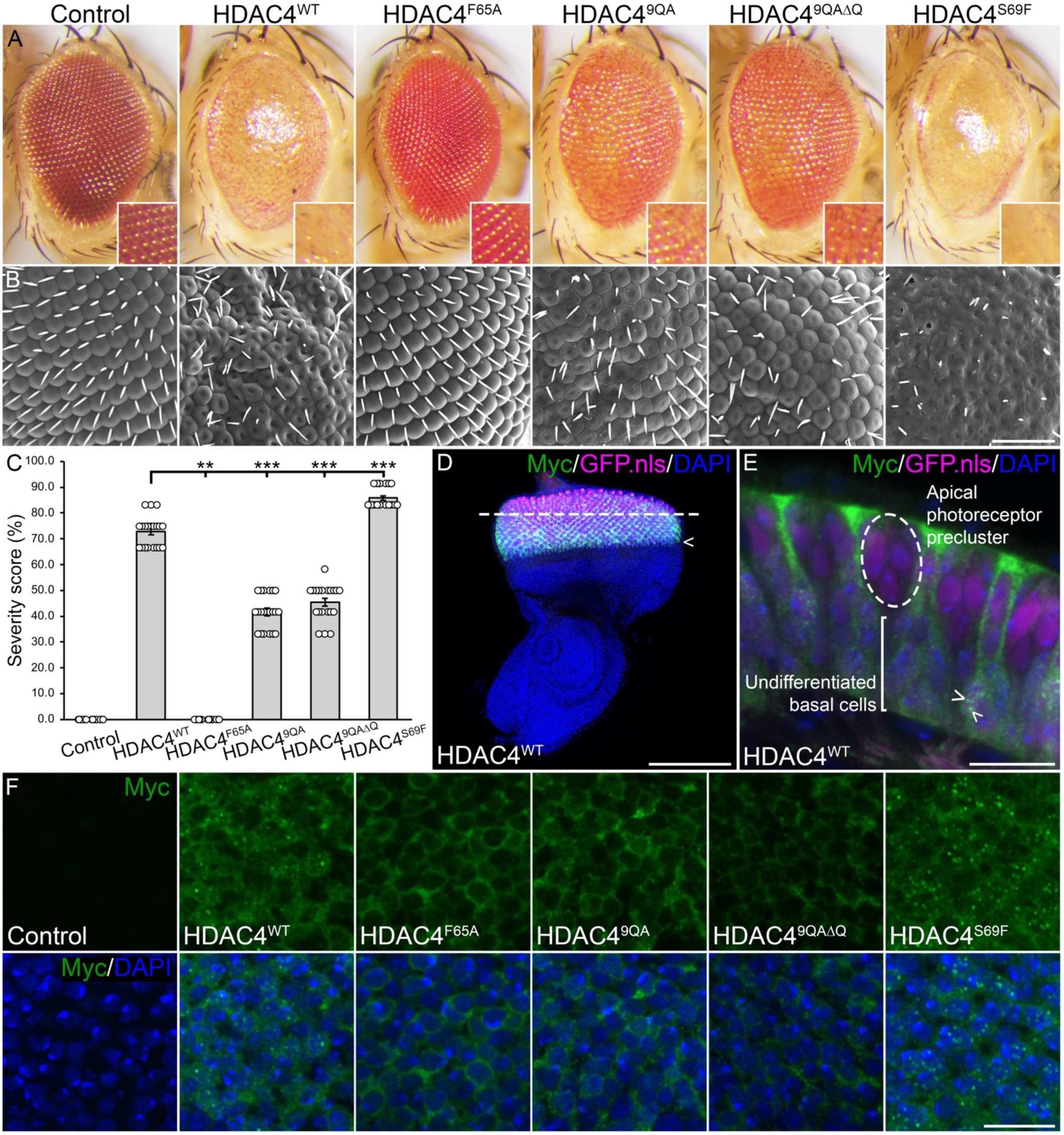
Oligomerization mutants alter HDAC4-overexpression induced defects in eye development. (A) Stereomicrographs (110x magnification) and (B) scanning electron micrographs of adult *Drosophila* eyes of flies raised at 25 °C. Flies carry one copy of *GMR-GAL4* and two copies of the *HDAC4-Myc* wild-type or mutant transgene (*GMR-GAL4/+; HDAC4/HDAC4*). The control is *GMR-GAL4/+; +.* Scale bar = 50 µm. (C) Quantification of eye phenotype severity. HDAC4^F65A^, HDAC4^9QA^ and HDAC4^9QAΔQ^ mutants were significantly reduced, and HDAC4^S69F^ was significantly increased in severity compared to HDAC4^WT^. ANOVA, F_(5,112)_ = 1642.19, p < 0.001, post-hoc Tukey’s HSD, *** p <0.001. Error bars indicate SEM. (D) Maximum projection of a third instar larval eye disc expressing HDAC4^WT^ (green) and GFP.nls (magenta) under the control of *GMR-GAL4*. Flies were raised at 25 °C. Expression of both HDAC4^WT^ and GFP.nls was observed posterior to the morphogenetic furrow (arrowhead). DAPI (blue) highlights nuclei across the whole disc. The dashed line indicates the location of the cross section depicted in (E). Scale bar = 100 µm. (E) Cross section of the larval eye disc demonstrating that HDAC4^WT^ localizes to both the apical and basal cells of the disc, but only forms small nuclear condensates (arrowheads) in undifferentiated basal cells. A photoreceptor precluster that forms a single ommatidium is circled in the apical surface, where HDAC4 condensates were not observed. Scale bar = 10 µm. (F) Single optical sections (0.5 µm) of basal cells of the larval eye disc expressing HDAC4^Mutant^-Myc (green), co-labelled with DAPI (blue). Scale bar = 10 µm.

### Oligomerization of HDAC4 regulates nuclear import

HDAC4 undergoes nucleocytoplasmic shuttling in response to stimuli (Chawla *et al*., 2003; Chen *et al*., 2014; Litke *et al*., 2018; Sando *et al*., 2012; Schlumm *et al*., 2013), with subcellular distribution differing between cell-types within and across species (Darcy *et al*., 2010; Main *et al*., 2021). Despite intrinsic nuclear import and export signals, nuclear import of HDAC4 is primarily mediated through MEF2 binding, with point mutations that abrogate MEF2 binding resulting in cytoplasmic retention (Wang & Yang, 2001). Previous studies showed that introducing the Phe93Asp mutation into hsHDAC4 (equivalent to DmHDAC4^F65A^) impaired MEF2-dependent transcriptional repression (Dai *et al*., 2024; Guo *et al*., 2007) suggesting that HDAC4 oligomerization may be necessary for MEF2 binding and proper subcellular localization.

To explore this hypothesis, HDAC4 distribution was examined in the *Drosophila* adult brain using *elav-GAL4*, which drives broader expression that *OK107-GAL4* (Hawley *et al*., 2023), allowing for the examination of HDAC4 subcellular distribution in cell-types additional to Kenyon cells. *elav-GAL4* also drives a considerably lower level of transgene expression in the mushroom body than *OK107-GAL4* (Hawley *et al*., 2023), and this is modest relative to the total levels of endogenous HDAC4 in the brain (Tan *et al*., 2024). This allows for the examination of how overexpression dose contributes to the regulation of HDAC4 subcellular distribution. Within Kenyon cell nuclei, HDAC4^WT^-Myc displayed a relatively uniform distribution (Fig. 6C,D), forming condensates that were smaller than those observed under *OK107-GAL4* expression (compared to Fig. 3), consistent with lower expression levels. Strikingly, HDAC4^F65A^, HDAC4^9QA^ and HDAC4^9QAΔQ^ displayed minimal nuclear localization, appearing as cytoplasmic haloes around the nuclei, whereas HDAC4^S96F^ formed larger and more numerous nuclear condensates than HDAC4^WT^.

**Figure 6.**
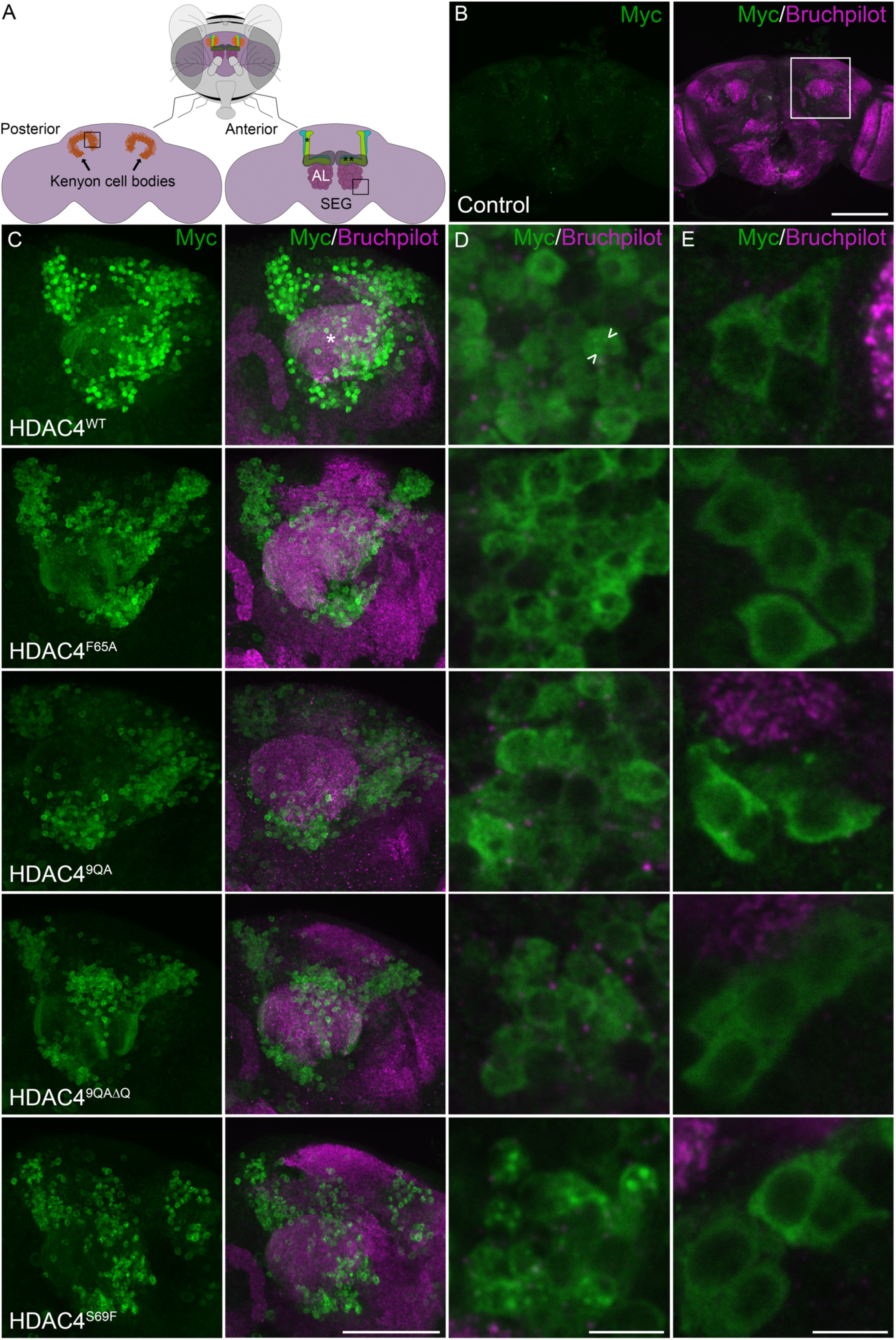
Expression and subcellular localization of HDAC4 oligomerization mutants in the adult brain. All genotypes were generated by crossing *elav-GAL4* females to males carrying the indicated *UAS-HDAC4-Myc* transgene and to the *w(CS10)* control. Whole mount brains were subjected to immunohistochemistry with anti-Myc (green) and anti-Bruchpilot (magenta), to highlight the neuropil structure in the brain. (A) Schematic showing the orientation of the brain in an adult fly (top), with the Kenyon cell bodies best viewed from the posterior (left) and the mushroom body lobes (* α, ** β/γ), antennal lobes (AL) and subesophageal ganglion (SEG) viewed from the anterior (right). Boxed region in the posterior brain is shown in (D), and in the anterior brain is shown in (E). (B) Maximum projection through the posterior of the brain. Boxed region is shown in (C). Scale bar = 100 µm. (C) Maximum projection through the calyx and Kenyon cell layer of the posterior of the brain. Asterisk denotes the calyx. Scale bar = 50 µm. (D-E) Magnified single optical section (0.5 µm) of HDAC4 in (D) Kenyon cells above the calyx and (E) cells surrounding the antennal lobe and subesophageal ganglion. Arrowheads point to nuclear condensates of HDAC4. Scale bars = 5 µm.

In cells surrounding the antennal lobes and the suboesophageal ganglion, HDAC4^WT^ was predominantly cytoplasmic, forming perinuclear haloes (Fig. 6E) and this distribution was unchanged for any of the mutants. Notably, these cells express a much lower level of MEF2 than Kenyon cells and MEF2 did not localize within HDAC4 nuclear condensates (SFig. 6). Together these data demonstrate that HDAC4 subcellular distribution varies by neuronal cell type, and is closely associated with MEF2 expression, supporting a role for oligomerization in nuclear entry and MEF2 binding.

### Oligomerization of HDAC4 in nuclei is required for HDAC4-induced disruption to development independent of its role in nuclear import

To assess the impact of nuclear retention on condensate formation, we generated HDAC4 mutants with compromised nuclear export, ensuring that phenotypic assessment was independent of differences in subcellular distribution. Specifically, we introduced mutations disrupting 14-3-3-dependent export, generating HDAC4^3SA-F65A^ and HDAC4^3SA-9QA^ (Tan *et al*., 2024; Wang *et al*., 2000; Wang & Yang, 2001). We hypothesized that the nuclear confinement of HDAC4^3SA-F65A^ would promote condensate formation through increasing local protein concentration. Since MEF2 is exclusively nuclear and its binding to HDAC4 promotes condensation and HDAC4-induced deficits (Tan *et al*., 2024), we generated HDAC4^3SA-F65A-ΔMEF2^, and also examined HDAC4^3SA-ΔMEF2^ (Tan *et al*., 2024) to test whether disrupting MEF2 binding would destabilize condensates. Expression was confirmed via western blot (Fig. 7A,B). To examine HDAC4 mutants in a MEF2-enriched background, they were expressed in Kenyon cells using *OK107-GAL4* in the presence of GAL80ts to avoid developmental epects. Fractionation assays showed that HDAC4^ΔMEF2^, HDAC4^9QA^ and HDAC4^F65A^ had reduced nuclear localization compared to HDAC4^WT^ (Fig. 7C-E), consistent withimmunohistochemistry (Fig. 6, Tan *et al*. (2024)), thus indicating that oligomerization supports nuclear entry. Incorporation of the 3SA mutation increased the nuclear distribution considerably, and this was not markedly reduced in any of the mutants (Fig. 7E), thus confirming their nuclear accumulation and enabling comparison of nuclear condensation without distribution bias.

**Figure 7.**
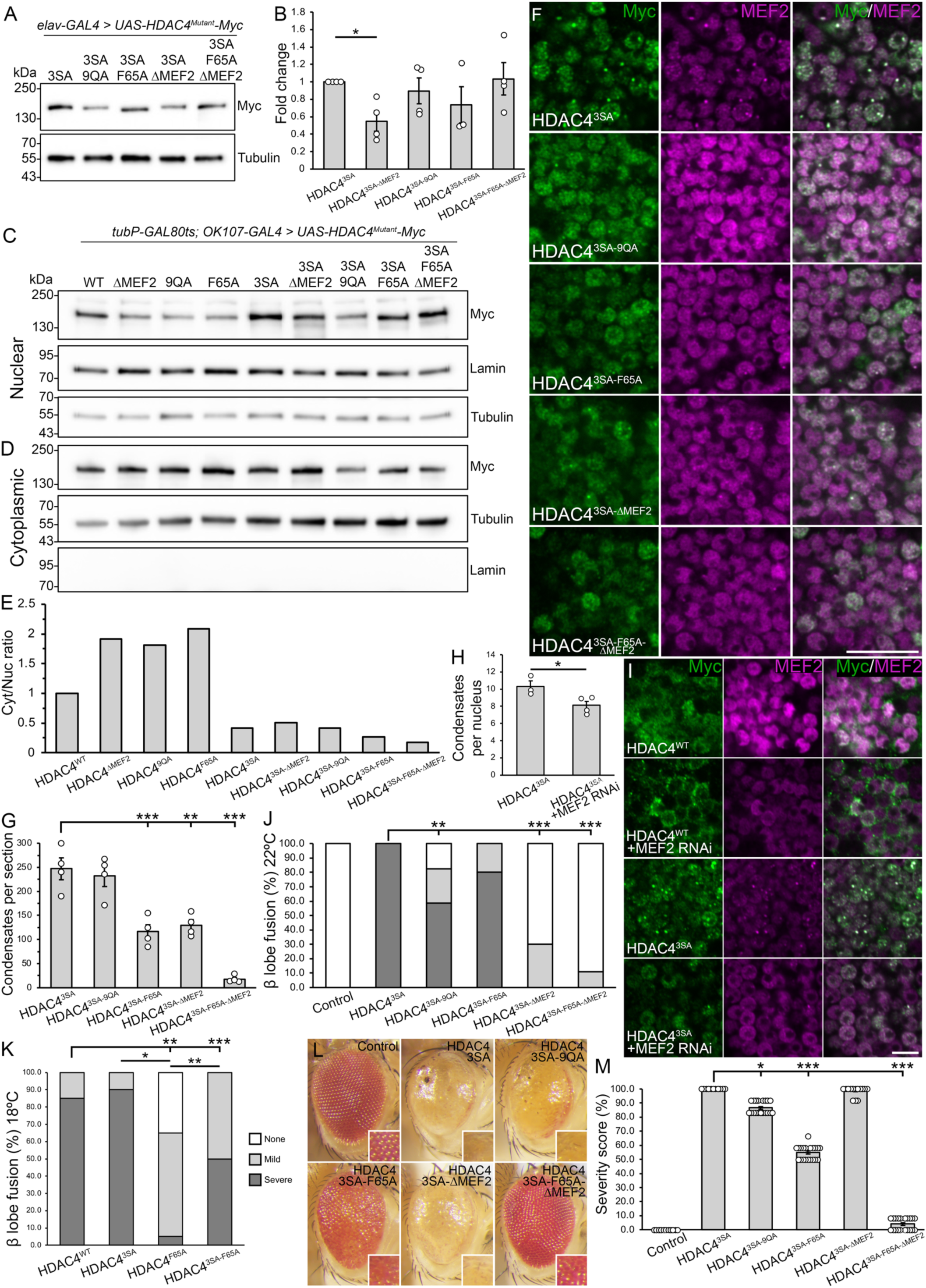
Functional analysis of nuclear HDAC4 oligomerization mutants and the role of MEF2. (A) Whole cell lysates generated from adult heads expressing *HDAC4^WT^-Myc* or *HDAC4^3SA-Mutant^-Myc* under the control of *elav-GAL4* were subjected to SDS-PAGE and probed for Myc and tubulin. (B) Quantification of HDAC4 band intensity normalized to tubulin (n = 4 blots). HDAC4^3SA-9QA^ and HDAC4^3SA-ΔMEF2^ were significantly reduced compared to HDAC4^3SA^. One sample t-test compared to HDAC4^3SA^; HDAC4^3SA-9QA^ t_(3)_ = 3.9078, p = 0.0298; HDAC4^3SA-F65A^ t_(3)_ = 0.7172, p = 0.525; HDAC4^3SA-ΔMEF2^ t_(2)_ = 1.1155, p = 0.3807; HDAC4^3SA-F65A-ΔMEF2^ t_(3)_ = 0.1755, p = 0.8719. Error bars indicate SEM. (C-D) Genotypes were generated by crossing *tubP-GAL80ts; OK107-GAL4* females to males carrying each indicated *UAS-HDAC4^Mutant^-Myc* transgene and to the *w(CS10)* control. Flies were raised at 18 °C until eclosion, and adults were transferred to 30 °C for three days to induce expression. Subcellular fractionation and western blotting was performed on fly heads. Membranes were probed with anti-Myc, as well as anti-lamin and anti-α-tubulin to assess nuclear and cytoplasmic abundance respectively, in each lysate. No lamin was detected in the cytoplasmic fractions. (E) Nuclear and cytoplasmic HDAC4 were quantified and normalized to lamin and tubulin respectively, before calculating the cytoplasmic to nuclear ratio relative to HDAC4^WT^. (F) Whole mount brains were subjected to immunohistochemistry with anti-Myc (green) and anti-MEF2 (magenta). Flies were raised as in (C). Representative single optical sections (0.5 µm) depict changes in condensate formation of nuclear-restricted HDAC4 in Kenyon cell nuclei. Scale bar = 10 µm. (G) Quantification of nuclear condensation of HDAC4 (performed as per Fig. 2D). ANOVA, F_(4,15)_ = 32.65, p < 0.0001; post-hoc Tukey’s HSD, ** p < 0.01, *** p < 0.001. Error bars indicate SEM. (H) Quantification of HDAC4^3SA^ condensates in the absence and presence of MEF2 RNAi (shown in I). Flies were raised as in (C) with males additionally carrying the UAS-MEF2-RNAi construct. The number of condensates (colocalizing puncta of HDAC4 and MEF2) per nucleus were counted and averaged for n ≥ 43 nuclei per section for n ≥ 3 brains per genotype. Unpaired t-test, HDAC4^3SA^ vs HDAC4^3SA^ + MEF2 RNAi t_(5)_ = 2.759, p = 0.0399. Error bars indicate SEM. (I) Whole mount brains were subjected to immunohistochemistry with anti-Myc (green) and anti-MEF2 (magenta). The distribution of HDAC4^WT^ shifted toward being more cytoplasmic in the presence of MEF2 knockdown while HDAC4^3SA^ remained nuclear but condensate number was decreased. (J-K) Quantification of β lobe fusion resulting from HDAC4 overexpression. Flies were raised at 22 (J) and 18 °C (K). HDAC4^3SA-9QA^, HDAC4^3SA-ΔMEF2^, and HDAC4^3SA-F65A-ΔMEF2^ mutants all displayed a significant reduction in severe β lobe fusion compared to HDAC4^3SA^ when expressed at 22 °C, p = 0.0019, p < 0.0001, and p < 0.0001 respectively. * p < 0.05, ** p < 0.01, *** p < 0.001, Fisher’s exact test. There was no significant difference in severe fusion between HDAC4^3SA^ and HDAC4^3SA-F65A^ (p = 0.1131) at 22 °C, but when expression was decreased by raising flies at 18 °C, a significant decrease was uncovered (p = 0.0138). (L) Stereomicrographs (110x magnification) of adult *Drosophila* eyes of flies raised at 25 °C. Flies carry one copy of *GMR-GAL4* and two copies of the *HDAC4^Mutant^-Myc* transgene (*GMR-GAL4/+; HDAC4/HDAC4*). The control is *GMR-GAL4/+; +.* (M) Quantification of eye phenotype severity. Significant differences between HDAC4^3SA^ and HDAC4^3SA-Mutant^ were observed. Kruskal-Wallis test, H_(5)_ = 106.92, p < 0.001, post-hoc Dunn’s test, * p < 0.05, *** p < 0.001. HDAC4^3SA^ vs HDAC4^3SA-9QA^, p = 0.01561; vs HDAC4^3SA-F65A^, p < 0.001; vs HDAC4^3SA-ΔMEF2^, p = 0.07861; vs HDAC4^3SA-F65A-ΔMEF2^ p < 0.001. HDAC4^3SA-F65A^ vs HDAC4^3SA-F65A-ΔMEF2^ p = 0.01387. Control vs HDAC4^3SA-F65A-ΔMEF2^ p = 0.3387). Error bars indicate SEM.

Condensate analysis revealed that HDAC4^3SA-9QA^ formed smaller condensates without significantly reducing their number (Fig. 7F,G). HDAC4^3SA-F65A^ produced smaller and fewer condensates, indicating that oligomerization supports condensate formation. When both oligomerization and MEF2 binding were disrupted, expression of HDAC4^3SA-F65A-ΔMEF2^ resulted in a synergistic decrease in the number of condensates. To confirm the role of MEF2 in condensate stabilization, we knocked down MEF2 via RNAi. This resulted in increased cytoplasmic localization of HDAC4^WT^ (Fig. 7I), as a consequence of reduced MEF2-dependent nuclear import. In contrast, since nuclear exit of HDAC4^3SA^ was blocked, this mutant remained confined to the nucleus, however in the presence of reduced levels of MEF2, condensates were smaller and fewer (Fig. 7H). Collectively these findings provide further support that MEF2 is a crucial determinant of nuclear HDAC4 condensate stability.

We next examined how these mutations apected mushroom body development. Expression of HDAC4^3SA^ caused severe β lobe fusion in 100% of brains, while HDAC4^3SA-9QA^ reduced this phenotype to 60% (Table 3, Fig. 7J). HDAC4^3SA-F65A^ expression decreased severe β lobe fusion to 80%, which was not significantly diperent from HDAC4^3SA^. Since HDAC4^3SA^ expression already caused maximal β lobe fusion, we lowered the temperature to reduce expression (Fig. 7K). Under these conditions, severe fusion was observed in 90% of HDAC4^3SA^ brains, but dropped to 50% with HDAC4^3SA-F65A^ expression. Strikingly, HDAC4^3SA-ΔMEF2^ and HDAC4^3SA-F65A-ΔMEF2^ mutations eliminated severe fusion (Table 3, Fig. 7J), and total fusion was reduced to 30% and 11% respectively. Together these data demonstrate that mutations impairing HDAC4 oligomerization reduce β lobe fusion, independent of subcellular distribution.

**Table 3.** Nuclear-restricted HDAC4 oligomerization mutants reduce HDAC4-overexpression induced defects in mushroom body development. All genotypes were generated by crossing *elav-GAL4* females to males carrying each *UAS-HDAC4-Myc* transgene and to the *w(CS10)* control. The percentage of brains displaying each phenotype was calculated from the total number of brains analysed for each genotype (n). At 22 **°**C overexpression of HDAC4^3SA^ resulted in a significant number of brains with β lobe fusion compared to control (p < 0.0001, Fisher’s exact test). HDAC4^3SA-9QA^, HDAC4^3SA-ΔMEF2^, and HDAC4^3SA-F65A-ΔMEF2^ mutants each significantly reduced severe β lobe fusion compared to HDAC4^3SA^ (p = 0.0019, p < 0.0001, and p < 0.0001 respectively) but there was no significant reduction by HDAC4^3SA-F65A^ (p = 0.1131, severe fusion). At 18°C severe β lobe fusion was significantly reduced for HDAC4^3SA-F65A^ compared to HDAC4^3SA^ (p = 0.0138) but HDAC4^3SA-F65A^ severity was increased compared to HDAC4^F65A^ (p = 0.0033). Severe β lobe fusion was reduced for HDAC4^F65A^ and HDAC4^3SA-F65A^ compared to HDAC4^WT^ (p < 0.0001 and p = 0.0407 respectively). Percentages may not sum to 100 due to rounding.

| 22 °C |  |  |  |  |  |  | 18 °C |  |  |  |  |
| --- | --- | --- | --- | --- | --- | --- | --- | --- | --- | --- | --- |
| (% of brains) | Control | HDAC4<sup>3SA</sup> | HDAC4<sup>3SA-9QA</sup> | HDAC4<sup>3SA-F65A</sup> | HDAC4<sup>3SA-ΔMEF2</sup> | HDAC4<sup>3SA-F65A-ΔMEF2</sup> | Control | HDAC4<sup>WT</sup> | HDAC4<sup>3SA</sup> | HDAC4<sup>F65A</sup> | HDAC4<sup>3SA-F65A</sup> |
| α lobe thinning | 0 | 50 | 27 | 35 | 5 | 0 | 0 | 10 | 40 | 0 | 10 |
| α lobe missing | 0 | 6 | 24 | 0 | 0 | 0.0 | 5 | 0 | 5 | 0 | 0 |
| β lobe thinning | 0 | 13 | 9 | 5 | 0 | 6 | 0 | 0 | 10 | 0 | 15 |
| β lobe missing | 0 | 13 | 0 | 5 | 5 | 11 | 0 | 0 | 5 | 10 | 5 |
| β lobe fusion | 0 | 100 | 82 | 100 | 30 | 11 | 20 | 100 | 100 | 65 | 100 |
| Mild | 0 | 0 | 24 | 20 | 30 | 11 | 20 | 15 | 10 | 60 | 50 |
| Severe | 0 | 100 | 59 | 80 | 0 | 0 | 0 | 85 | 90 | 5 | 50 |
| Guidance defect | 0 | 50 | 32 | 45 | 10 | 11 | 5 | 0 | 50 | 15 | 40 |
| No defects | 100 | 0 | 6 | 0 | 60 | 78 | 75 | 0 | 0 | 25 | 0 |
| n | 20 | 16 | 34 | 20 | 20 | 18 | 20 | 20 | 20 | 20 | 20 |

Expression of HDAC4^3SA^ also severely disrupted eye development, resulting in complete fusion of ommatidia, loss of pigmentation, and loss of bristles (Fig. 7L,M). Eyes also appeared shrunken toward the ventral side, and necrosis was observed. Expression of HDAC4^3SA-9QA^ slightly reduced the severity with fewer necrotic spots, though fusion and loss of pigmentation remained. HDAC4^3SA-F65A^ partially improved the phenotype, allowing identification of some individual ommatidia. Pigmentation was also increased, but fusion and disorganization persisted. Notably, HDAC4^3SA-F65A-ΔMEF2^ almost fully rescued the phenotype, with eyes appearing largely wild-type with normal pigmentation and no fusion. In contrast, HDAC4^3SA-ΔMEF2^ produced only a minor improvement compared to HDAC4^3SA^. It is therefore evident that Phe65 is crucial for nuclear HDAC4-induced disruption to eye development, while the MEF2 binding site acts synergistically with Phe65 to exacerbate the phenotype. Since we previously illustrated that MEF2 binding is not required by HDAC4 to disrupt eye development, and detected no endogenous expression of MEF2 in the developing eye (Tan *et al*., 2024), these findings suggest an additional protein responsible in part for the eye phenotype may be present in photoreceptor condensates, the interaction of which is mediated at the MEF2 binding site.

## Discussion

Here we demonstrate that oligomerization of HDAC4 is essential for nuclear condensation and contributes to overexpression-induced neurodevelopmental defects in *Drosophila*. By using mutants with disrupted oligomerization or MEF2 binding, we dissected the roles of these HDAC4 functional regions in condensate formation and neurotoxicity. We determined that oligomerization promotes condensate formation, as HDAC4^F65A^ and HDAC4^9QA^ reduced both oligomerization and condensate number, and this decrease correlated with reduced neurodevelopmental defects in both the mushroom body and adult eye. However, the direct correlation between nuclear condensates and phenotype severity was somewhat confounded, as these mutations also reduced the nuclear abundance of HDAC4. To address this, we enforced nuclear localization of these mutants by introducing the 3SA mutation. Despite this, HDAC4^3SA-F65A^ and HDAC4^3SA-9QA^ still reduced condensate size and number, confirming that oligomerization contributes to nuclear condensation. Moreover, loss of MEF2 binding further destabilized condensates. Consistent with this, the strongest phenotypic rescue was observed when Phe65 was substituted in addition to disrupting MEF2 binding. This underscores a cooperative role for oligomerization and MEF2 binding in HDAC4 nuclear accumulation and subsequent condensate formation. Together, these findings establish HDAC4 oligomerization and MEF2 binding as key drivers of HDAC4-induced neurotoxicity, linking nuclear condensates to neurodevelopmental pathology.

The most common disruption to HDAC4 observed in neuronal disease and disease models is an abnormal increase in its nuclear concentration. HDAC4 is a dynamic protein with many binding partners that modulate its function, including its movement between the nucleus and cytoplasm. The most well characterised interaction is with MEF2, which colocalizes in nuclear condensates (Fitzsimons *et al*., 2013). The interaction between MEF2 and HDAC4 is complex; HDAC4 is responsible for the repression of MEF2-dependent transcription (Miska *et al*., 1999; Sparrow *et al*., 1999; Wang *et al*., 1999), while MEF2 mediates nuclear import of HDAC4 (Borghi *et al*., 2001; Wang & Yang, 2001). Increased abundance of MEF2 increases HDAC4 nuclear localization and condensation (Tan *et al*., 2024), while mutation of the MEF2-binding domain or knockdown of MEF2 reduced nuclear condensation independent of changes in oligomerization and subcellular distribution respectively, demonstrating that MEF2 not only increases nuclear localization but stabilises condensation of HDAC4. It is therefore conceivable that changes in MEF2 may result in altered nucleocytoplasmic shuttling of HDAC4 that induce nuclear accumulation in neuronal disease.

Protein aggregation is a highly contentious subject in the literature, especially when arising in the context of an overexpression study as the physiological significance can be unclear. However, it is evident that both oligomerization and condensation of HDAC4 has biological relevance with respect to its wild-type function, as well as in a disease context. HDAC4 is observed in small granular or punctate foci in the nucleus and cytoplasm of neurons in non-disease mouse and *Drosophila* brains, as well as along axons and dendrites (Darcy *et al*., 2010; Tan *et al*., 2024). We hypothesize that these granular accumulations represent wild-type HDAC4 function, acting as potential regulatory sites of transcription factors such as MEF2. Such sites are highly dynamic and likely chromatin-bound, given the presence of MEF2. However, upon increased HDAC4 nuclear abundance, regulation of the formation of these sites is disrupted, resulting in their coalescence into larger condensates. Indeed, nuclear accumulation of HDAC4 is observed in some Alzheimer’s disease brains, and the degree of accumulation positively correlated with Braak staging (Herrup *et al*., 2013; Shen *et al*., 2016). The correlation between nuclear condensation of HDAC4 and severity of neurodevelopmental phenotypes in *Drosophila* therefore supports that these condensates may be mediating neuronal dysfunction in a disease context.

An additional important consideration in the context of neuronal disease is the dominant negative effect of HDAC4 condensates. We previously demonstrated that endogenous HDAC4 is sequestered in condensates that result from GAL4-induced expression of transgene HDAC4 (Tan *et al*., 2024), and here we show that endogenous HDAC4 likely supports the oligomerization and condensation of the HDAC4 mutants that are themselves defective in oligomerization. Wakeling *et al*. (2021) identified seven heterozygous HDAC4 missense mutations in unrelated individuals, all affecting a 14-3-3 binding site. Functional analysis of two of the mutations confirmed reduce 14-3-3 binding *in vitro*, which were predicted to promote nuclear localization and retention of HDAC4 *in vivo*. Given the dominant negative nature of HDAC4 condensation, the mutant pool of HDAC4 in these patients likely drives condensation of wild-type HDAC4, exacerbating mutation-associated phenotypes despite heterozygosity. These mutations have further significance in light of the recent finding that 14-3-3 binding to HDAC4 destabilises condensate formation (Liu *et al*., 2024). Further elucidating the role of 14-3-3 in HDAC4 condensate dynamics will therefore be of significant interest as a potential therapeutic strategy to reduce HDAC4 nuclear accumulation and condensation.

Oligomerization of HDAC4 regulates the association of MEF2 and thus nucleocytoplasmic shuttling of HDAC4. We show that the mechanism of HDAC4 oligomerization is conserved between human and *Drosophila* HDAC4; mutations that disrupted the hydrophobic core or stabilising polar interaction networks prevented N-terminal oligomerization *in vitro* and reduced full-length oligomerization and condensation *in vivo*. The sequences that mediate these interactions are not only conserved among species, but also within the class IIa HDACs (Guo *et al*., 2007), suggesting that oligomerization is essential for wild-type class IIa HDAC function. Indeed, oligomerization is required for HDAC4-dependent repression of MEF2-dependent transcription *in vitro* (Dai *et al*., 2024; Guo *et al*., 2007), and we observed mutations that disrupted oligomerization of HDAC4 increased its cytoplasmic distribution in Kenyon cells. In support of a functional relationship between HDAC4 oligomerization and MEF2 binding, Dai *et al*. (2024) recently demonstrated that when the N-terminus of hsHDAC4 is bound by the HDAC-interacting domain of MEF2, HDAC4 forms a dimer that facilitates chromatin looping via the formation of a hydrophobic core involving Phe93, identical to that which mediates tetramerization (Guo *et al*., 2007). These data highlight how HDAC4 oligomerization facilitates not only MEF2 binding and nuclear import, but the formation of regulatory sites via the recruitment of transcription factors. Therefore, examining the chromatin binding profile of HDAC4 and how this changes with altered nuclear abundance and condensation will be of interest in determining mechanisms of neurodisruption.

We have clearly demonstrated that HDAC4 nuclear condensate formation is dependent on dose, and given that increased nuclear abundance of HDAC4 is observed in neuronal disease, it is likely that increased levels and/or increased nuclear localization of HDAC4 precedes visible condensation. HDAC4 condensates are not β-sheet insoluble inclusions, but instead are highly dynamic and display features of liquid-liquid phase separated (LLPS) droplets in that their formation is both concentration and interaction-dependent. Phase separation involves a nucleation even that seeds condensate formation (Alberti & Dormann, 2019). The protein that nucleates LLPS is often termed a scaffold, and this interacts with client proteins that partition into condensates following seeding. It is possible that HDAC4 oligomerization may constitute this nucleation event, and that MEF2 acts as a client and further stabilises condensation. Another key feature of LLPS is valency, which describes how the scaffold protein has many binding regions and motifs, including intrinsically disordered regions (IDRs) (Nam & Gwon, 2023). Indeed regions outside of the N-terminus of human HDAC4 are predicted to be intrinsically disordered with the potential for mediating phase separation (Liu *et al*., 2024), and we demonstrated these are required for condensation of *Drosophila* HDAC4. Importantly, together this suggests that smaller puncta may too undergo phase separation, further supporting the idea that they act as novel sites for regulation of interacting factors.

Glutamine-rich regions are frequently implicated in protein aggregation (Chen *et al*., 2001). Interestingly however, we have demonstrated that glutamine residues are essential to HDAC4 protein stability. While deleting the significant polyglutamine stretch in DmHDAC4 did not further destabilise condensation, we observed reduced condensate size and/or number for each of the HDAC4^9QA^, HDAC4^9QAΔQ^, and HDAC4^3SA-9QA^ mutants *in vivo*, and they were also reduced in protein levels. Similarly, *in vitro* the purified HDAC4^N-9QA^ was difficult to handle, rapidly degraded, and did not appear to form an α-helix. However, it still appeared to retain the ability to oligomerize. Given the dose dependence of HDAC4 condensate formation, we therefore surmise that the reduced condensation observed *in vivo* was a function of reduced protein levels and not solely decreased oligomerization. Moreover, the reduced dose of each of the glutamine mutants prevented us from effectively assessing the contribution of reduced oligomerization by this mechanism to overexpression-induced phenotypes as HDAC4 overexpression-induced phenotypes in the eye and mushroom body are sensitive to dose (Schwartz *et al*., 2016; Tan *et al*., 2024). Interestingly, mutating only seven of the glutamine residues within an N-terminal construct of human HDAC4 did not affect its stability (Guo *et al*., 2015), suggesting that each may contribute to N-terminal and full-length HDAC4 stability to a different degree. Understanding the residues important in HDAC4 turnover will be essential as this could be used to modulate levels of HDAC4 to prevent nuclear accumulation and condensation.

We have used *Drosophila* as a model of neuronal development previously to understand the different roles of nuclear and cytoplasmic HDAC4 (Main *et al*., 2021; Tan *et al*., 2024) and here to specifically understand the effects of HDAC4 condensation on neuronal function. This model has effectively highlighted the subtleties in function of different neuronal subtypes, and thus the different effects HDAC4 condensate formation has on each. While HDAC4 overexpression and condensation disrupts both mushroom body and eye development, we previously demonstrated that an intact MEF2 binding site is required for manifestation of the mushroom body but not the eye phenotype (Tan *et al*., 2024). Interestingly, here we observed that mutation of the MEF2 binding site in HDAC4^3SA-F65A-ΔMEF2^ significantly reduced eye phenotype severity compared to HDAC4^3SA-F65A^. Despite its presence in the optic lobes of the brain (Schulz *et al*., 1996) we have not been able to detect endogenous expression of MEF2 within developing photoreceptor nuclei (Tan *et al*., 2024) suggesting the mechanism through which HDAC4 affects eye development is MEF2-independent. The MEF2 binding site of human HDAC4 overlaps with the binding site of a number of proteins, including that for serum response factor (SRF) (Backs *et al*., 2011; Wang & Yang, 2001), the *Drosophila* orthologue of which is implicated in rhabdomere development (Gambis *et al*., 2011) and may thus be implicated in HDAC4 condensate formation as well as overexpression induced defects in eye development. It is therefore imperative that the cell type in which HDAC4 condensation is observed is considered. MEF2 isoform expression varies throughout development and in adult brains, often with non-overlapping patterns (reviewed in Lisek *et al*. (2023)), thus it is essential to consider both MEF2-dependent and independent mechanisms through which HDAC4 may disrupt neuronal function.

## Conclusion

In summary, HDAC4 condensate formation occurs in a highly dynamic manner, dependent on oligomerization, dose, subcellular distribution, and protein-protein interactions. We have demonstrated a correlation between nuclear condensation of HDAC4 and defects in neurodevelopment in *Drosophila*, suggesting HDAC4 condensates may mediate neuronal dysfunction in neurodevelopmental and neurodegenerative disease. These condensates contain not only HDAC4, but also MEF2, and we hypothesise additional binding partners are present, therefore phenotypes are mediated by a combination of altered HDAC4 function as well as altered function of proteins sequestered in these condensates. MEF2 is a key mediator of the mushroom body phenotype, with its HDAC4 interaction disrupted by altered oligomerization. However, HDAC4 binding partners important in the eye phenotype have not yet been identified, demonstrating that HDAC4 function differs between neuronal populations. These findings have relevance for understanding both wild-type HDAC4 function with respect to the role of oligomerization, and highlight that HDAC4 may be a therapeutic target for diseases in which HDAC4 regulation is disrupted.

## Methods

### Fly strains

All flies were maintained on standard medium at 22 °C with a 12-hour light dark cycle unless otherwise indicated. *P{w[+mW.hs]=GawB}elav[c155]* (*elav-GAL4*, BDSC #458), *w[^∗^]; P{w[+mW.hs]=GawB}OK107 ey[OK107]/In(4)ci[D], ci[D] pan[ciD] sv[spa-pol]* (*OK107-GAL4*, BDSC #854), *w[^∗^]; P{w[+mC]=GAL4-ninaE.GMR}12* (*GMR-GAL4*, BDSC #1104) and *w[1118]; P{w[+mC]=UAS-GFP.nls}14* (*UAS-GFP.nls*, BDSC #4775) and P{w[+mC]=UAS-APP.Abeta42.B}m26a (*UAS-Aβ42*, BDSC #33770) were obtained from the Bloomington *Drosophila* Stock Center. *w^∗^; P{w+mC = tubP-GAL80ts}10; TM2/TM6B, Tb1* (*tubP-GAL80ts*) and *w(CS10)* strains were provided by R. Davis (The Scripps Research Institute, Jupiter, FL). P{GD5039}v15550 (UAS-*MEF2-RNAi*, VDRC #15550) was obtained from the Vienna *Drosophila* Resource Center. *GFP-HDAC4^WT^* was previously generated (Main *et al*., 2021). Homozygous lines carrying *OK107-GAL4*, *tubP-GAL80ts,* and both *GFP-HDAC4^WT^* and *HDAC4^Mutant^-Myc* were generated by standard genetic crosses. The open reading frame of wild-type *Dm HDAC4* was previously synthesized (nucleotides 461–4216, NCBI NM_132640) with a C-terminal 6x Myc tag (Main *et al*., 2021), *HDAC4^WT^-Myc*, and mutagenesis performed by GenScript (New Jersey, United States) to obtain *HDAC4^F65A^, HDAC4^9QA^, HDAC4^9QAΔQ^, HDAC4^S69F^, HDAC4^3SA-9QA^, HDAC4^3SA-F65A^, HDAC4^3SA-F65A-ΔMEF2^, as well as HDAC4^3SA^ and HDAC4^3SA-ΔMEF2^* as previously described (Main *et al*., 2021). Replacement of the *HDAC4^WT^* 6x Myc tag with a 3x HA tag was carried out by Genscript. *HDAC4^M1-L285^-HA* (nucleotides 461– 1315, NCBI NM_132640), with the Ser239Ala mutation was synthesized with a C-terminal 3x HA tag. *HDAC4* constructs were cloned into the pUASTattB vector for germline transformation of *Drosophila* (Genetivision, Houston, TX) using the P2 docking site at (3L)68A4. All strains were outcrossed for a minimum of five generations into the *w(CS10)* genetic background.

### Protein expression, purification, and biochemical analyses

The hsHDAC4 tetramer structure (RCSB PDB (RRID:SCR_012820) entry 2H8N, Guo *et al*. (2007)) was examined and represented using UCSF ChimeraX (Meng *et al*. (2023), https://www.cgl.ucsf.edu/chimerax/, version 1.7, RRID:SCR_015872). Protein sequence alignment of *Drosophila* (nucleotides 461–4216, NCBI NM_132640, translated) and human (NCBI NP_006028) HDAC4 was performed using EMBOSS Water pairwise sequence alignment (Madeira *et al*. (2024), https://www.ebi.ac.uk/jdispatcher/psa/emboss_water, RRID:SCR_025141). Residues Ala62-Gln153 of hsHDAC4 were previously examined for tetramer formation (Guo *et al*., 2007) and the corresponding region of DmHDAC4 was determined as Pro37-Gln143. This region was codon optimized for *E. coli* and synthesized by GenScript to create HDAC4^N-WT^. This was followed by mutagenesis to create HDAC4^N-F65A^, HDAC4^N-9QA^ and HDAC4^N-S69F^. HDAC4^N^ constructs were cloned into the pET-15B expression vector in frame with the N-terminal 6x His tag. HDAC4^N^ were transformed into BL21(DE3) *E. coli* for recombinant protein expression. Cells were grown at 30 °C in 2x YT culture medium containing ampicillin (100 µg/mL) until OD_550_ reached ∼0.6, at which time expression was induced with 0.5 mM IPTG and cells were grown at 20 °C overnight (∼14 hours, HDAC4^WT^ and HDAC4^S69F^, 1 L total culture volume) or at 25 °C for three hours (HDAC4^F65A^ and HDAC4^9QA^, 10 L total culture volume). Cells were harvested by centrifugation at 5,000 x g for 10 minutes and pellets stored at -80 °C. Pellets were thawed and resuspended in 20 mL (per L of culture) lysis buffer (30 mM HEPES, pH 7.5, 300 mM NaCl, 10 µg/mL Pepstatin A, 1X cOmplete EDTA-free protease inhibitors (Roche)) before lysis via Cell Disruptor (Constant Systems) at 10 kpsi and sonication for 2x 30 second pulses at 20 W. The lysate was clarified by centrifugation at 20,000 x g for 30 minutes before filter sterilization. The supernatant was subjected to nickel-affinity chromatography using Profinity IMAC Resin (BioRad) in IMAC buffer (30 mM HEPES, pH 7.5, 300 mM NaCl) with a gradient elution from 30 mM-500 mM imidazole. A_215_ and A_230_ were examined to determine the fractions in which HDAC4^N^ eluted, and these were also run on SDS-PAGE to confirm HDAC4^N^ presence given the low number of tyrosine and tryptophan residues. Pooled fractions containing HDAC4^N^ were subjected to size exclusion chromatography on a Superdex 75 10/300 GL column (Cytiva) in SEC buffer (30 mM HEPES, pH 7.6, 150 mM NaCl), fractions containing HDAC4^N^ pooled and concentrated to ∼0.5-2 mg/mL before snap freezing and storing at -80 °C.

Crosslinking was performed as per Guo *et al*. (2007). Briefly, 0.5 mg/mL HDAC4^N^ diluted in SEC buffer (20 µL) was incubated with 2 µL disuccinimidyl suberate (DSS, Thermo Fisher Scientific) dissolved in DMSO for 30 minutes at room temperature. The reaction was quenched for 15 minutes with 1 µL of 1M Tris (pH 7.5) before 5 µL was prepared for SDS-PAGE followed by colloidal Coomassie staining. Circular dichroism was performed on HDAC4^N^ prepared in SEC buffer at 0.5 mg/mL in a 0.1 mm quartz cuvette. Spectra were recorded using a Chirascan Spectrophotometer (Applied Photophysics). For each sample spectra were collected between 180-260 nm in 1 nm steps and averaged, and data were baseline corrected for the sample buffer.

### SDS-PAGE & western blotting

Whole cell lysates were produced by homogenizing snap-frozen *Drosophila* heads in either IP buffer (50mM sodium chloride, 30 mM sodium pyrophosphate, 50 mM sodium fluoride, 10% glycerol, 0.5% Triton X-100, 0.5mM PMSF, 25mM Tris, pH 7.05, 1X protease inhibitors (cOmplete EDTA free Protease Inhibitor Cocktail (Roche)) or RIPA buffer (150 mM sodium chloride, 0.1% Triton X-100, 0.5% sodium deoxycholate, 0.1% SDS, 50 mM Tris pH 8.0, 1X protease inhibitors) and collecting the supernatant following centrifugation at 12,000 x g for two minutes. Nuclear and cytoplasmic lysates were prepared using the NE-PER nuclear and cytoplasmic extraction reagents (Thermo Fisher Scientific) using a protocol modified from Maitra *et al*. (2019). Heads (n = 50) were homogenized in 100 µL of cytoplasmic buffer 1 (CB1) supplemented with protease inhibitors and incubated on ice for 10 minutes before adding 5.5 µL of cytoplasmic buffer 2 (CB2) and centrifugation at 10,000 x g for five minutes. The supernatant was retained as the cytoplasmic fraction, and the pellet washed twice by resuspending in 50 µL of CB1, incubating for 10 minutes, then adding 2.75 µL of CB2, centrifuging at 10,000 x g for five minutes and discarding the supernatant. The pellet was resuspended in 50 µL of nuclear extraction reagent with protease inhibitors, incubated on ice for 40 minutes and centrifuged at 10,000 x g for 10 minutes before collecting the supernatant as the nuclear fraction. Total protein was quantified using the Pierce BCA Protein Assay kit (Thermo Fisher Scientific). Whole cell lysate (30 µg) or subcellular fractions (20 µg) were denatured in 1X sample buffer (2% SDS, 5% 2-mercaptoethanol, 10% glycerol, 0.01% bromophenol blue, 60 mM Tris HCl pH 6.8) at 90 °C for five minutes. Samples were loaded onto 4-20% or 10% Mini-PROTEAN TGX Precast Protein gels (BioRad) and electrophoresed at 200 V. Protein was blotted onto a nitrocellulose membrane for one hour at 4°C before blocking in 5% skim milk powder in TBST (20 mM Tris, 150 mM NaCl, pH 7.6, 0.1% Tween-20) for one hour at room temperature. Following washing in TBST membranes were incubated overnight at 4°C in primary antibody in 1% skim milk powder/TBST, and one hour at room temperature in secondary antibody. Antibodies used were rabbit anti-Myc (ab9106, Abcam, 1:1,000, Antibody Registry Identifier RRID:AB_307014), rabbit anti-GFP (ab290, Abcam, 1:4,000, RRID:AB_303395), rat anti-HA (Roche, clone 3F10, 1:1,000, RRID:AB_2687407), mouse anti-α-Tubulin (12G10 clone, Developmental Studies Hybridoma Bank (DSHB), 1:500, RRID:AB_1157911), mouse anti-lamin(Dm0) (ADL67.10 clone, DSHB, RRID:AB_528336), sheep anti-mouse-HRP (Sigma Aldrich NA931VS, 1:20,000), donkey anti-rabbit-HRP (Sigma Aldrich NA934VS, 1:40,000), goat anti-rat-HRP (Abcam ab97057, 1:10,000, RRID:AB_10680316). Amersham ECL Prime detection reagent (GE Life Sciences) was used as per manufacturer’s instructions, and chemiluminescence detected using the Azure Biosystems C600 imaging system. Quantification of western blots was performed using the Gel Analyzer plugin within ImageJ (Schneider *et al*., 2012) (https://imagej.net/, RRID:SCR_003070). Whole cell and cytoplasmic lysate signal were normalized to tubulin, and nuclear lysate to lamin. For blots of whole cell lysates, the fold-change of the normalized signal was calculated relative to HDAC4^WT^. For blots from subcellular fractionation, the nuclear to cytoplasmic ratio was calculated for each sample and normalized to HDAC4^WT^.

### Immunoprecipitation

Protein A/G magnetic beads (Thermo Fisher Scientific, 25 µL) were resuspended and washed in TBS-T (20 mM Tris, 150 mM NaCl + 0.05% Tween-202) before incubation with 2 µL anti-GFP (ab290, Abcam, RRID:AB_303395) for one hour at room temperature. Whole cell lysate (2 mg) was added to the antibody-bead mixture and incubated overnight at 4°C. Beads were washed three times in TBS-T and proteins eluted in 1X sample buffer for 10 minutes at room temperature. The supernatant was collected and incubated at 95 °C for five minutes. Total eluate was loaded for SDS-PAGE and western blotting.

### Immunohistochemistry

Whole flies were prefixed in PFAT-DMSO (4% paraformaldehyde, PBS, 0.1% Triton X-100, 5% DMSO) for one hour at room temperature, followed by washing and dissection in PBT (10 mM phosphate buffer, pH 7.4, 0.5% Triton X-100). For mushroom body analyses, adult flies were anethetized with CO_2_ and placed on ice prior to dissection. Third instar crawling larvae were washed and the dissected in PBS (10 mM phosphate buffer, pH 7.4). Tissues were post-fixed in PFAT-DMSO for 20 minutes and stored in 100% methanol. Tissues were rehydrated in 50% methanol/PBT, blocked for three hours at room temperature in immunobuffer (5% normal goat serum in PBT) and incubated overnight at room temperature with primary antibody (mouse anti-Bruchpilot, 1:100, DSHB clone nc82, RRID:AB_2314866; mouse anti-FasII, 1:20, DSHB clone 1D4, RRID:AB_528235; rabbit anti-GFP, 1:20,000, Abcam ab290, RRID:AB_303395; rat anti-HA, 1:500, Roche clone 3F10 RRID:AB_2687407; rabbit anti-MEF2, 1:500, gift from Bruce Paterson, National Cancer Institute, Bethesda; rabbit anti-Myc, 1:100, Abcam ab9106, RRID:AB_307014; mouse anti-Myc, 1:100, DSHB clone 9E10, RRID:AB_2266850), followed by overnight incubation at 4 °C with secondary antibody (goat anti-mouse Alexa 555, 1:500, Invitrogen A-21422, RRID:AB_2535844; goat anti-rabbit Alexa 647, 1:500, Invitrogen A21244, RRID:AB_2535812; goat anti-rat Alexa 555, 1:500, Invitrogen A21434, RRID:AB_2535855), then washing in PBT. Brains were incubated with FSB (0.01% in 50% ethanol) or THT (0.25% in 50% ethanol), washed in 50% ethanol, and then washed further in PBT. Samples that required nuclear staining were washed in PBT before incubation with DAPI (1:20,000 in PBS) and all tissues were mounted in Antifade (90% glycerol, 0.2% n-propyl gallate, 10 mM phosphate buffer, pH 7.4 (Sigma Aldrich), 0.5% Triton X-100). Images were captured using a Leica TCS SP5 DM6000B Confocal Microscope or a Zeiss LSM900 super-resolution microscope, and processed using ImageJ software.

### Live imaging

Live imaging of adult *Drosophila* brains was performed using a Zeiss LSM900 super-resolution microscope using a protocol modified from Savoian (2017). Brains were dissected from live flies in PBS and immediately mounted in Voltalef 10S oil, a coverslip placed atop, and imaged using a 63x oil immersion lens.

### Light microscopy

Light microscopy was performed using an Olympus SZX16 Stereo Zoom microscope and CellSens Dimension (Olympus) imaging software. Flies were frozen at -20°C before thawing and imaging at 110x magnification. Constant light intensity and exposure was used. Images were imported into Adobe Photoshop (RRID:SCR_014199), Z-axis drift accounted for using the Auto-Align Layers function, and optical sections stacked using the Auto-Blend Layers function.

### Scanning electron microscopy

Scanning electron microscopy was performed as previously described (Main *et al*., 2021). Briefly, adult flies were anaesthetized and immersed in primary modified Karnovsky’s fixative (3% glutaraldehyde, 2% formaldehyde, 0.1 M sodium phosphate buffer and Triton X-100), followed by vacuum infiltration, ethanol dehydration, and critical point drying. Heads were mounted onto aluminium stubs, sputter coated with gold (Bal-Tec SCD 050 sputter coater), and imaged using an FEI Quanta 200 environmental scanning electron microscope (20.00 kV, spot 4.0 nm).

### Quantification of condensation and neurodevelopmental phenotypes

HDAC4 and MEF2 condensates in Kenyon cell nuclei were counted using the Cell Counter plugin (https://imagej.net/ij/plugins/cell-counter.html, RRID:SCR_025376) in ImageJ and were only counted if they were visible in both single and merged channels. For panels in Figs. 2D, 7G and SFig. 4 condensates were counted through ten serial optical sections at 1 µm increments, beginning at the first emergence of the calyx (n = 4 brains per genotype), imaged at 150x magnification. When condensates were present in adjacent slices they were only counted once. For panels in Figs. 3I, K and 7H, condensates per nucleus were counted in a single optical section (705x magnification) for ≥ 27 cells per section (n ≥ 3 brains per genotype) using QuPath (Bankhead *et al*. (2017), https://qupath.github.io/, RRID:SCR_018257). Statistical significance was assessed using a one-way ANOVA with post-hoc Tukey’s HSD test with significance set at α = 0.05.

Assessment of mushroom body phenotypes was performed by a blinded scorer as per Tan *et al*. (2024). Z-stacks of mushroom body lobes were scored for the presence or absence of developmental defects in the α- and β lobes, including lobe thinning, absence, and guidance defects, as well as β lobe fusion. Fusion was scored as mild if axons measuring less than half the width of the lobe crossed the midline of the brain, or as severe if axons measuring more than half the width of the lobe crossed the midline. Statistical significance was assessed using the Fisher’s exact test.

Assessment of eye development was performed by a blinded scorer as per Tan *et al*. (2024). A scoring system was developed to assess severity of defects in morphology which considered three key facets of eye development that can be reproducibly analysed; bristle formation, ommatidia alignment, fusion and pigmentation. Each aspect was scored from 0-4, for a total severity score of 12 per eye, which was converted to % severity. Bristles: 0 = All bristles present and correctly placed, 1 = Most bristles correctly placed, with no more than a small number missing or extra, 2 = Moderate loss of bristles or misplaced bristles and/or large loss of bristles at edges of the eye, 3 = Few bristles, 4 = Complete absence of bristles. Ommatidia: 0 = Ommatidia correctly aligned and no fusion, 1 = Alignment of ommatidia altered but no fusion, 2 = Ommatidia alignment perturbed, altered depth of ommatidia boundary but no fusion, 3 = Ommatidia alignment perturbed and fusion, 4 = Complete absence of ommatidia (total fusion). Pigmentation: 0 = No changes in pigmentation, 1 = Small number of ommatidia missing pigment, 2 = Moderate loss of pigmentation across the eye, 3 = Pigmentation only around the edge of the eye, or in small spots across eye, 4 = No pigmentation. Where data were normally distributed statistical significance was assessed using a one-way ANOVA with post-hoc Tukey’s HSD test with significance set at α = 0.05. Where data were not normally distributed, the Kruskal Wallis test was used with significance set at α = 0.05, followed by a post-hoc Dunn’s test with a Bonferroni corrected α.

## Acknowledgements

We thank Bruce Paterson for the MEF2 antibody, and Yi-Hsuan (Jennifer) Tu & Trevor Loo for assistance with protein purification. We also thank the Manawatū Microscopy and Imaging Centre, Massey University for assistance with confocal microscopy.

