## Supplementary data for "N-terminal oligomerization drives HDAC4 nuclear condensation and neurodevelopmental dysfunction in *Drosophila*"

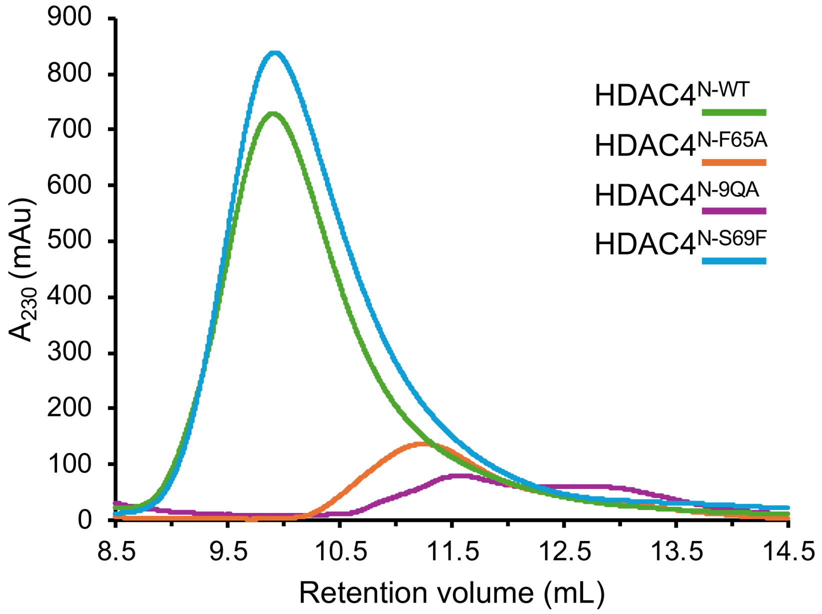


***Supplementary Figure 1. HDAC4$^N$ mutants vary in oligomerization as assessed by size exclusion chromatography.*** Size exclusion chromatography profiles of wild-type and mutant HDAC4$^N$ immediately after nickel-affinity chromatography purification. Pooled fractions from the affinity purification were directly injected onto a Superdex 75 10/300 column. Due to the lack of aromatic amino acids, absorbance was recorded at 230 nm as previously reported (Dai *et al.*, 2024).

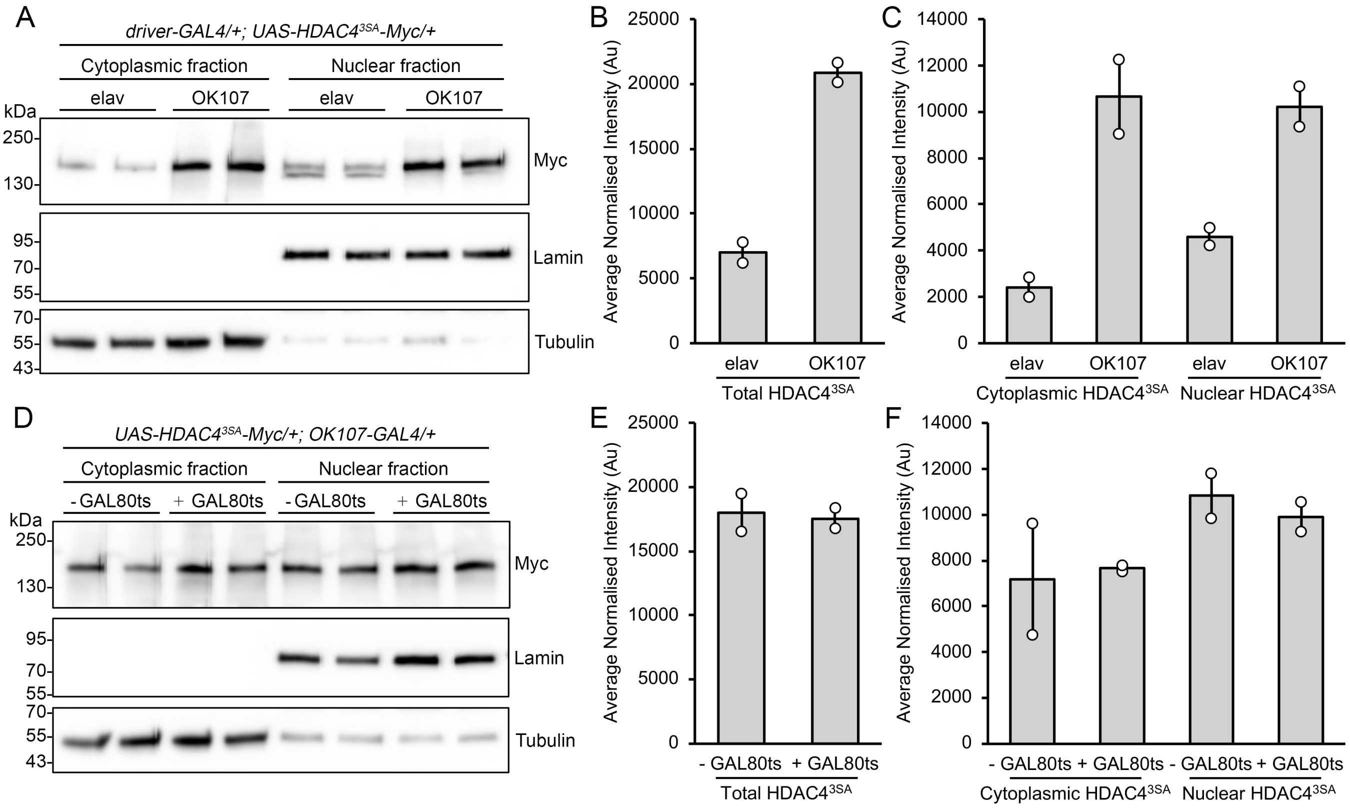


***Supplementary Figure 2. Quantification of HDAC4$^{3SA}$ in nuclear and cytoplasmic fractions under the control of elav-GAL4 and OK107-GAL4 in the absence and presence of tubP-GAL80ts.*** (A,D) Genotypes were generated by crossing *elav-GAL4* (elav), *OK107-GAL4* (OK107 -GAL80ts) or *tubP-GAL80ts; OK107-GAL4* (OK107 +GAL80ts) females to males carrying *UAS-HDAC4$^{3SA}$-Myc*. Flies were raised at 18 °C until eclosion, and adults were transferred to 30 °C for three days to induce expression. Subcellular fractionation and western blotting was performed on heads of flies. Membranes were probed with anti-Myc, as well as anti-lamin and anti-$\alpha$-tubulin to assess nuclear and cytoplasmic abundance respectively in each lysate. No lamin was detected in the cytoplasmic fractions. (B-C, E-F) Quantification of band intensity. Nuclear and cytoplasmic HDAC4$^{3SA}$ intensity was measured and normalized to lamin and tubulin respectively, then the normalized values were added (nuc + cyt) for each independent preparation and averaged between preparations to give average normalized intensity. (B) Total HDAC4$^{3SA}$ intensity was increased under the control of *OK107-GAL4* compared to *elav-GAL4*. (D) No differences in HDAC4$^{3SA}$ intensity were observed in the presence or absence of GAL80ts. (C,F) Quantification of nuclear and cytoplasmic band intensity. Nuclear and cytoplasmic HDAC4$^{3SA}$ intensity was measured, and normalized to lamin and tubulin respectively, then the normalized values were added for each nuclear and cytoplasmic preparation (nuc + nuc, cyt + cyt) and averaged to give average normalized intensity. (C) HDAC4$^{3SA}$ intensity for each cytoplasmic and nuclear fraction was increased for *OK107-GAL4* compared to *elav-GAL4*. (F) HDAC4$^{3SA}$ intensity for each cytoplasmic and nuclear fraction varied but there was no significant difference in the presence or absence of GAL80ts.

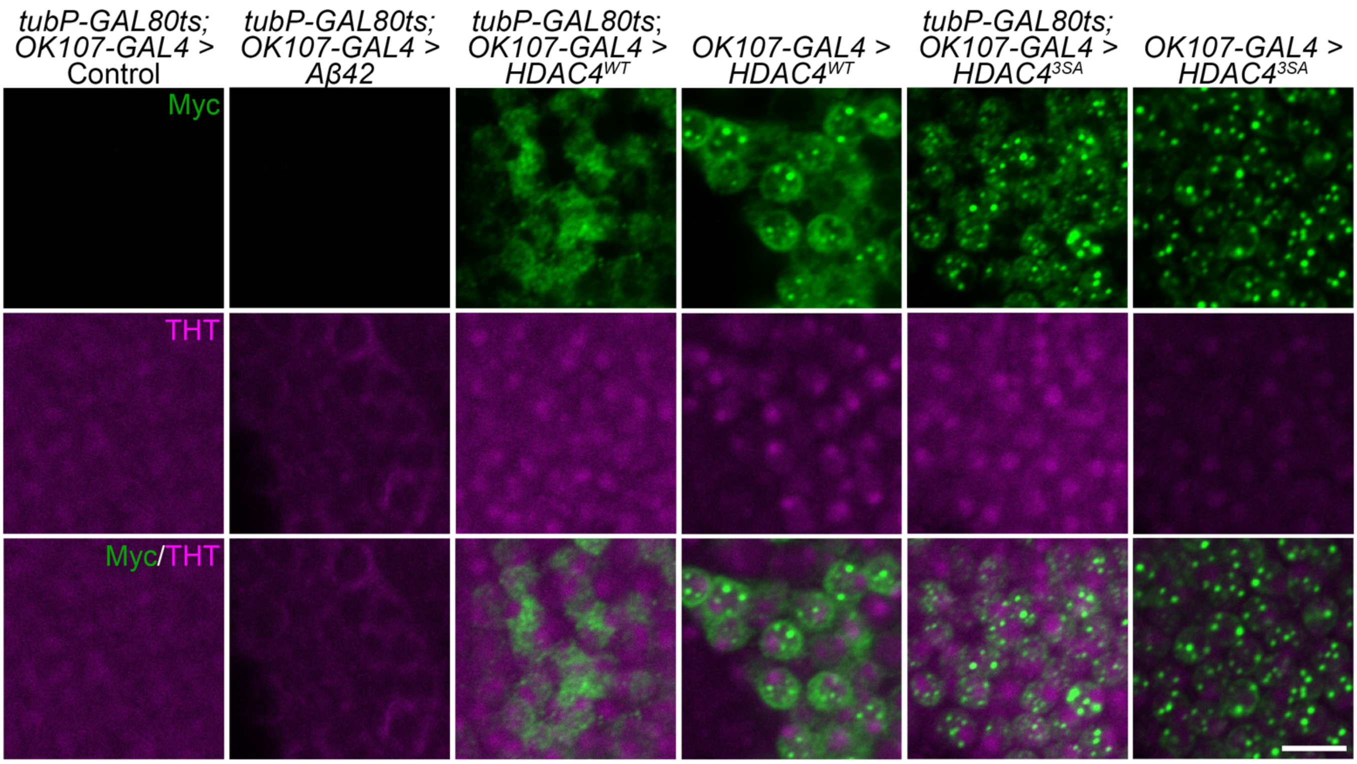


***Supplementary Figure 3. HDAC4 condensates do not colocalize with thioflavin T.*** Whole mount brains were subjected to immunohistochemistry with anti-Myc (green) and counterstained with thioflavin-T (THT, magenta). Representative single optical sections (0.5 µm) through the Kenyon cell layer are shown. Genotypes were generated by crossing *tubP-GAL80ts; OK107-GAL4,* or *OK107-GAL4* females to males carrying the noted UAS-transgene and the *wCS10* control, and flies were raised at either 18 °C throughout development and transgene expression was induced by incubation of flies at 30 °C for 72 hours (+ tubP-GAL80ts) or raised at 25 °C throughout development and adulthood (- tubP-GAL80ts). THT successfully detected Aβ42, but did not bind HDAC4 condensates.

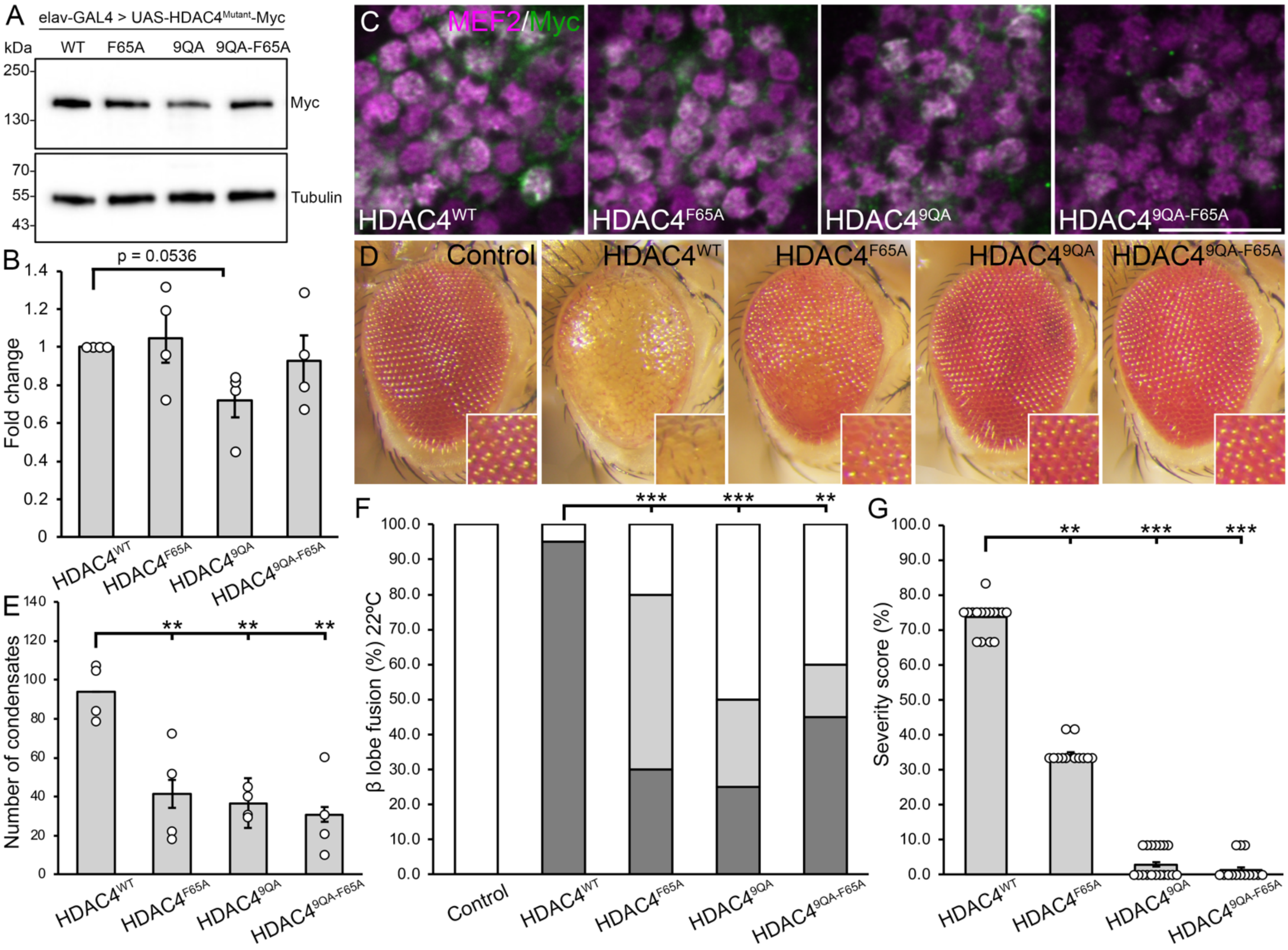


***Supplementary Figure 4. Combination of HDAC4$^{9QA}$ and HDAC4$^{F65A}$ mutations does not further reduce condensate formation or overexpression-induced phenotypes.*** (A) Whole cell lysates generated from adult heads *expressing HDAC4$^{WT}$-Myc* or *HDAC4$^{Mutant}$-Myc* under the control of *elav-GAL4* were subjected to SDS-PAGE and probed for Myc and tubulin. (B) Quantification of HDAC4 band intensity normalized to tubulin (n = 4 blots). HDAC4$^{9QA}$ was almost significantly reduced compared to HDAC4$^{WT}$, and there was no difference between each of HDAC4$^{F65A}$ and HDAC4$^{9QA\text{-}F65A}$ compared to HDAC4$^{WT}$. One sample t-test compared to HDAC4$^{WT}$, HDAC4$^{9QA}$ $t_{(3)} = 3.0915$, $p = 0.0536$; HDAC4$^{F65A}$ $t_{(3)} = 0.3717$, $p = 0.7348$; HDAC4$^{9QA\text{-}F65A}$ $t_{(3)} = 0.5502$, $p = 0.6205$. Error bars indicate the SEM. (C) Genotypes were generated by crossing *tubP-GAL80ts; OK107-GAL4* females to males carrying the indicated *UAS-HDAC4-Myc* transgene and transgene expression was induced in adulthood using GAL80ts. Flies were raised at 18 °C until eclosion when adults were transferred to 30 °C for 72 hours to induce transgene expression. Whole mount brains were subjected to immunohistochemistry with anti-Myc (green) and anti-MEF2 (magenta). Representative single optical sections (0.5 µm) through the calyx and Kenyon cell layer of the posterior of the brain showing HDAC4 condensates and colocalization with MEF2. Scale bar = 10 µm. (D) Stereomicrographs (110x magnification) of adult *Drosophila* eyes raised at 25 °C. Flies carry one copy *of GMR-GAL4* and two copies of the indicated *HDAC4-Myc* transgene (*GMR-GAL4/+; HDAC4/HDAC4*). Control is *GMR-GAL4/+ ;* +. (E) Quantification of nuclear condensation of HDAC4. The number of condensates (colocalizing puncta of HDAC4 and MEF2) per optical section were counted and averaged for 10 sections through the posterior of the brain at 1 µm increments (n = 4 brains per genotype). There was no significant difference between HDAC4$^{9QA\text{-}F65A}$ compared to each of HDAC4$^{9QA}$ and HDAC4$^{F65A}$ (ANOVA, $F_{(3,12)} = 9.7956$, $p = 0.00151$, post-hoc Tukey's HSD, $p = 0.8533$ and $p = 0.9716$ respectively. ** $p < 0.01$). (F) Quantification of β lobe fusion resulting from HDAC4 overexpression. Flies were raised at 22 °C. There was no statistically significant difference in severe β lobe fusion associated with expression of HDAC4$^{9QA\text{-}F65A}$ compared to each of HDAC4$^{9QA}$, $p = 0.5145$, and HDAC4$^{F65A}$, $p = 0.3203$, Fisher's exact test. (G) Quantification of the severity of eye phenotypes. There was no significant difference between HDAC4$^{F65A}$ and HDAC4$^{9QA\text{-}F65A}$, Kruskal-Wallis test, $H_{(3)} = 72.21$, $p < 0.001$, post-hoc Dunn's test, $p = 0.5719$. ** $p < 0.01$, *** $p < 0.001$. Error bars indicate SEM.

| (% of brains) | Control | HDAC4$^{WT}$ | HDAC4$^{9QA}$ | HDAC4$^{F65A}$ | HDAC4$^{9QA\text{-}F65A}$ |
|---|---|---|---|---|---|
| α lobe thinning | 0 | 35 | 0 | 10 | 0 |
| α lobe missing | 0 | 0 | 0 | 0 | 0 |
| β lobe thinning | 0 | 25 | 5 | 0 | 0 |
| β lobe missing | 0 | 0 | 5 | 5 | 5 |
| **β lobe fusion** | **0** | **95** | **80** | **50** | **60** |
| Mild | 0 | 0 | 50 | 25 | 15 |
| Severe | 0 | 95 | 30 | 25 | 45 |
| Guidance defect | 0 | 0 | 10 | 5 | 5 |
| No defects | 100 | 5 | 15 | 40 | 35 |
| n | 20 | 20 | 20 | 20 | 20 |

***Supplementary Table 1. Combination of 9QA and F65A mutations does not further reduce HDAC4-overexpression induced defects in mushroom body development.*** All genotypes were generated by crossing *elav-GAL4* females to males carrying the indicated *UAS-HDAC4-Myc* transgene and to the *w(CS10)* control. The percentage of brains displaying each phenotype was calculated from the total number of brains analysed for each genotype (n). Statistical analysis was performed with the Fisher's Exact Test. Overexpression of HDAC4$^{WT}$ resulted in a significant number of brains with β lobe fusion compared to control ($p < 0.0001$). There was no significant reduction in severe β lobe fusion for HDAC4$^{9QA\text{-}F65A}$ compared to HDAC4$^{9QA}$ ($p = 0.5145$) and HDAC4$^{F65A}$ ($p = 0.3203$).

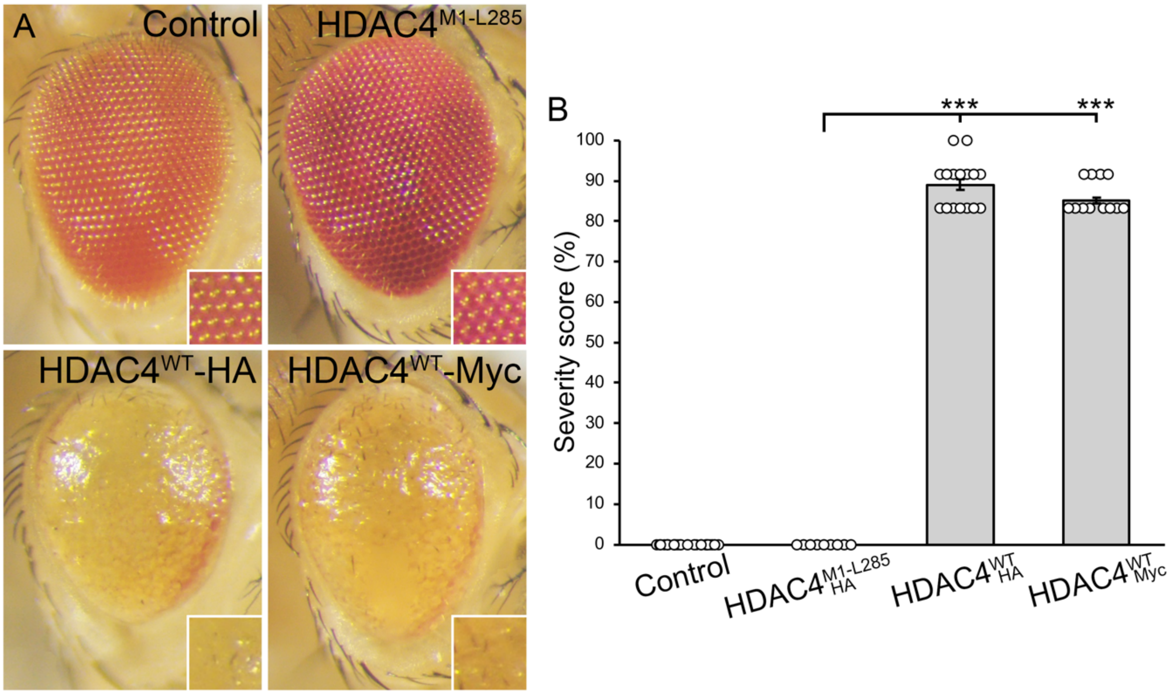


***Supplementary Figure 5. HDAC4$^{M1\text{-}L285}$ does not induce defects in eye development.*** (A) Stereomicrographs (110x magnification) of adult *Drosophila* eyes raised at 25 °C. Flies carry one copy of *GMR-GAL4* and two copies of the *HDAC4* transgene (*GMR-GAL4/+; HDAC4/HDAC4*). Control is *GMR-GAL4/+ ; +*. (B) Quantification of eye phenotype severity. HDAC4$^{WT}$-HA and HDAC4$^{WT}$-Myc induced a severe phenotype compared to controls, while none was observed for HDAC4$^{M1\text{-}L285}$. Kruskal-Wallis test, $H_{(3)} = 71.73$, $p < 0.001$, post-hoc Dunn's test, *** $p < 0.001$. Control vs HDAC4$^{M1\text{-}L285}$, $p = 1$. HDAC4$^{M1\text{-}L285}$ vs

$HDAC4^{WT}$-HA, $p < 0.001$; vs $HDAC4^{WT}$-Myc, $p < 0.001$. $HDAC4^{WT}$-HA vs $HDAC4^{WT}$-Myc, $p = 0.272$. Error bars indicate SEM.

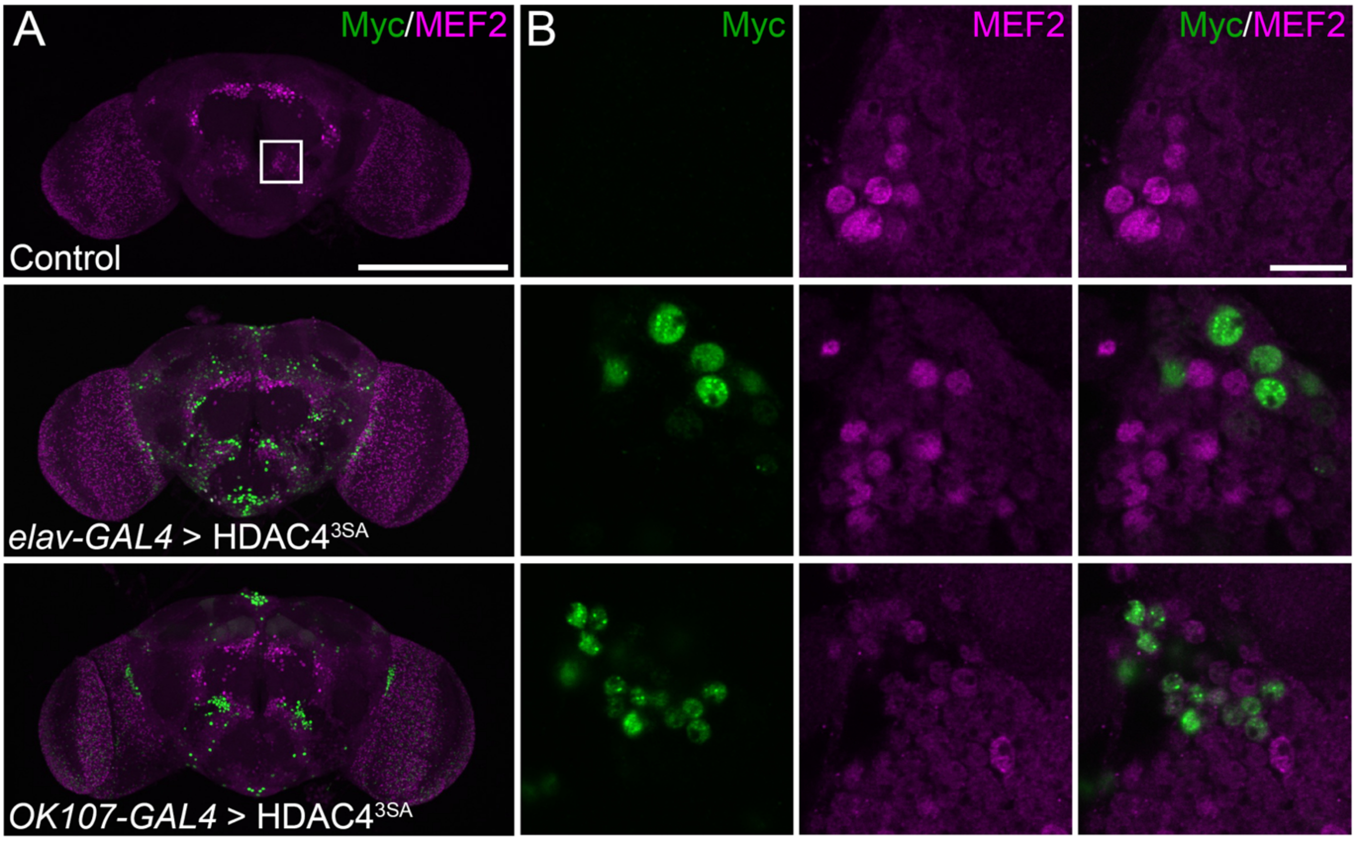


***Supplementary Figure 6. MEF2 levels are reduced in cells surrounding the antennal lobe and subesophageal ganglion.*** Whole mount brains were subjected to immunohistochemistry with anti-Myc (green) and anti-MEF2 (magenta). Genotypes were generated by crossing *elav-GAL4* and *OK107-GAL4* females to *$HDAC4^{3SA}$-Myc* males. The control was *wCS10* wild-type flies. Flies were raised at 18 °C until eclosion when adults were transferred to 30 °C for 72 hours to increase transgene expression. (A) Maximum projections through the anterior of the brain showing endogenous MEF2 expression in the presence of $HDAC4^{3SA}$ overexpression compared to the control. White boxed region is depicted in (B). Scale bar = 200 μm. (B) Single optical sections (0.5 μm) through nuclei below the antennal lobe. HDAC4 nuclear condensates formed, however these did not appear to sequester MEF2, and MEF2 levels were lower in these nuclei than those in surrounding regions in which HDAC4 was not expressed. Scale bar = 10 μm.

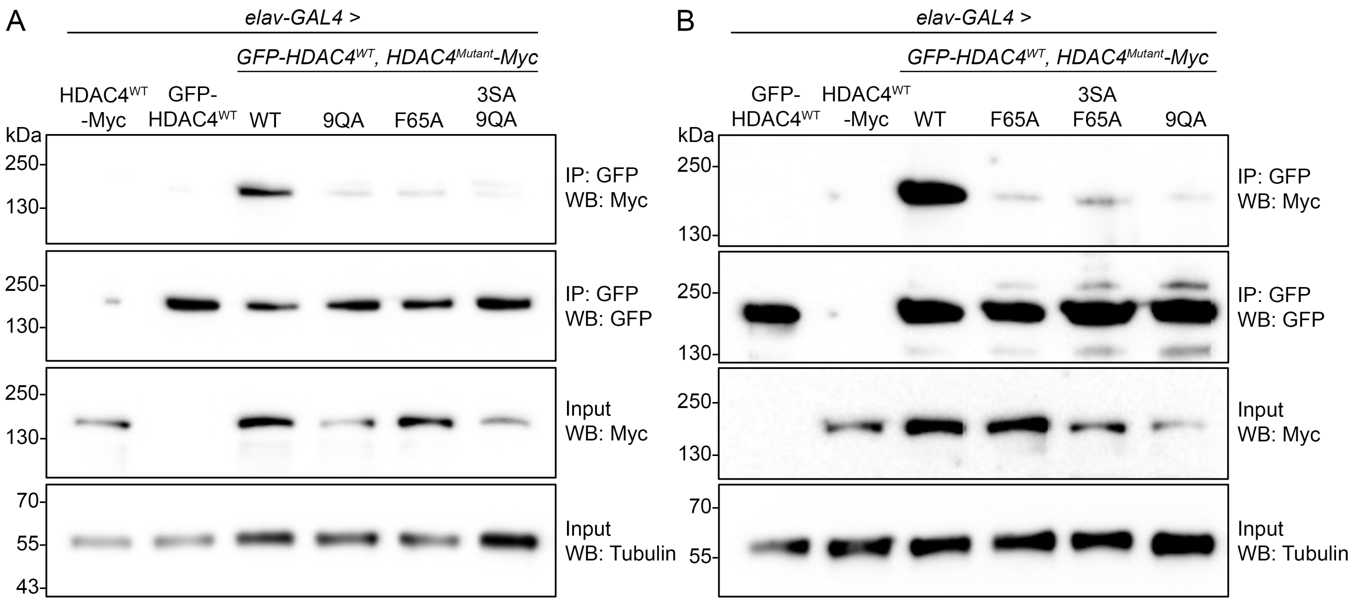


***Supplementary Figure 7. HDAC4$^{3SA\text{-}F65A}$ and HDAC4$^{3SA\text{-}9QA}$ exhibit a severely impaired oligomerization capacity.*** (A,B) Mutation of the hydrophobic core or conserved glutamine residues alters oligomerization of HDAC4$^{3SA}$ *in vivo*. Genotypes were generated by crossing *elav-GAL4* females to males carrying the indicated *HDAC4* transgene. Flies were raised at 18 °C until eclosion when adults were transferred to 22 °C to increase transgene expression. Co-immunoprecipitation was performed as described in Figure 1F,G. Co-immunoprecipitation yield of HDAC4$^{3SA\text{-}9QA}$ (A, lane 6) and HDAC4$^{3SA\text{-}F65A}$ (B, lane 5) is significantly reduced compared to HDAC4$^{WT}$ (lane 3) upon immunoprecipitation of GFP-HDAC4$^{WT}$. An alternative protocol was used to generate the data shown in (B); Protein A/G PLUS agarose beads were used, and whole cell lysates were first pre-cleared by incubation with unconjugated beads.

***Supplementary Figure 8. Full western blots and gels.***

**Fig. 1C**

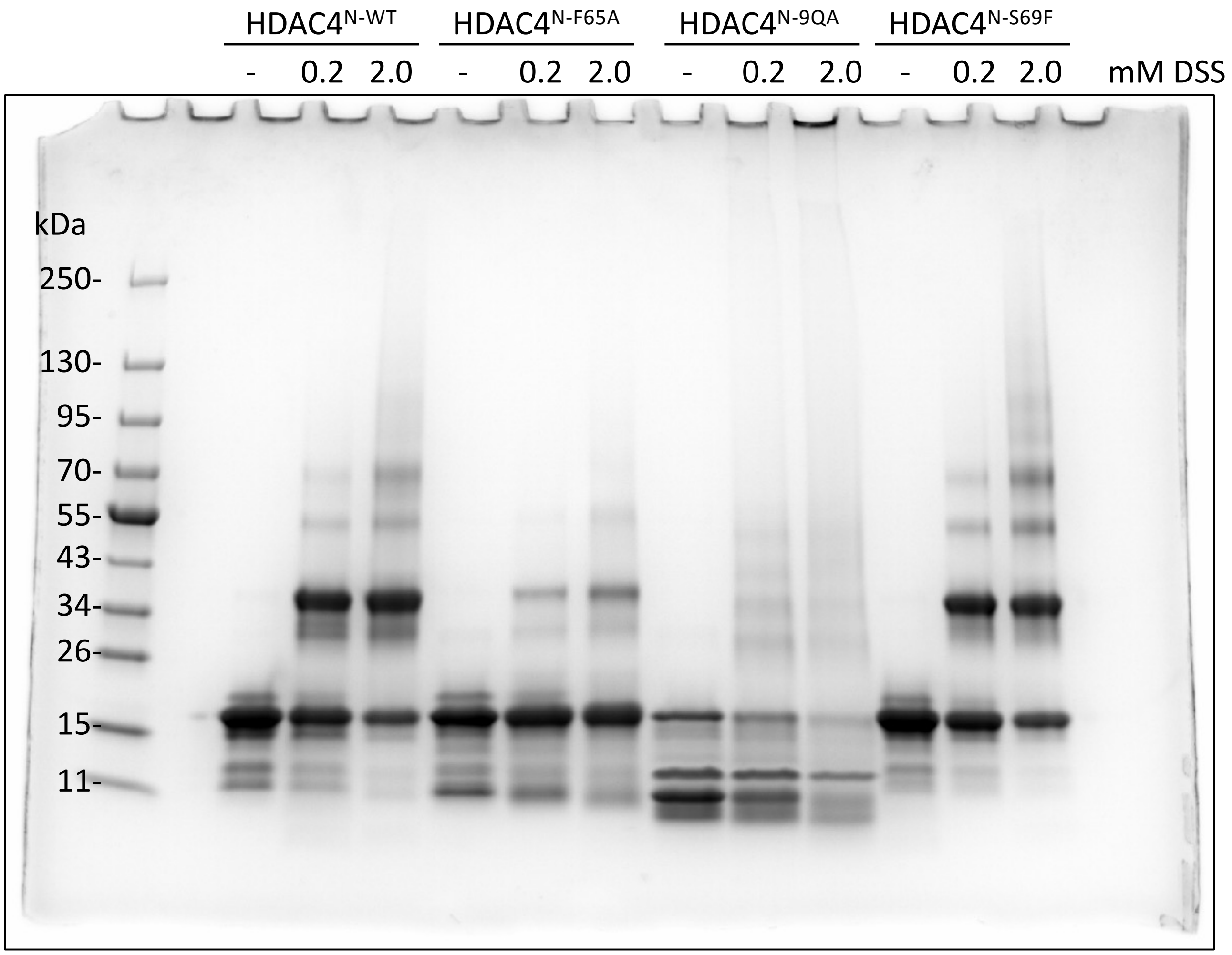
HDAC4$^{N-WT}$
HDAC4$^{N-F65A}$
HDAC4$^{N-9QA}$
HDAC4$^{N-S69F}$
- 0.2 2.0 - 0.2 2.0 - 0.2 2.0 - 0.2 2.0 mM DSS
kDa
250-
130-
95-
70-
55-
43-
34-
26-
15-
11-

**Fig. 1G**

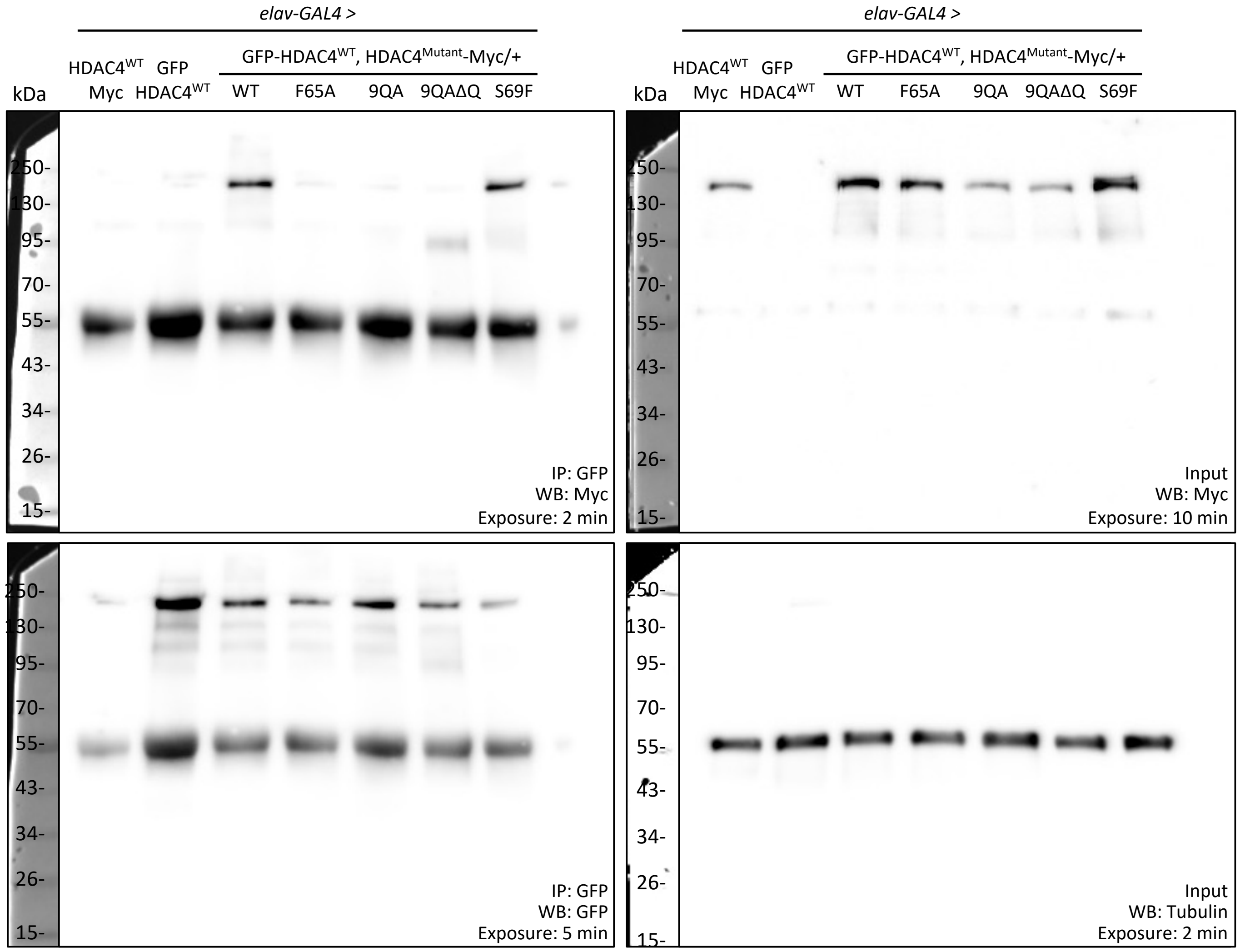


The band at ~50 kDa in the IP blots (left) corresponds to the IgG heavy chain detection.

**Fig. 1H & S4A**

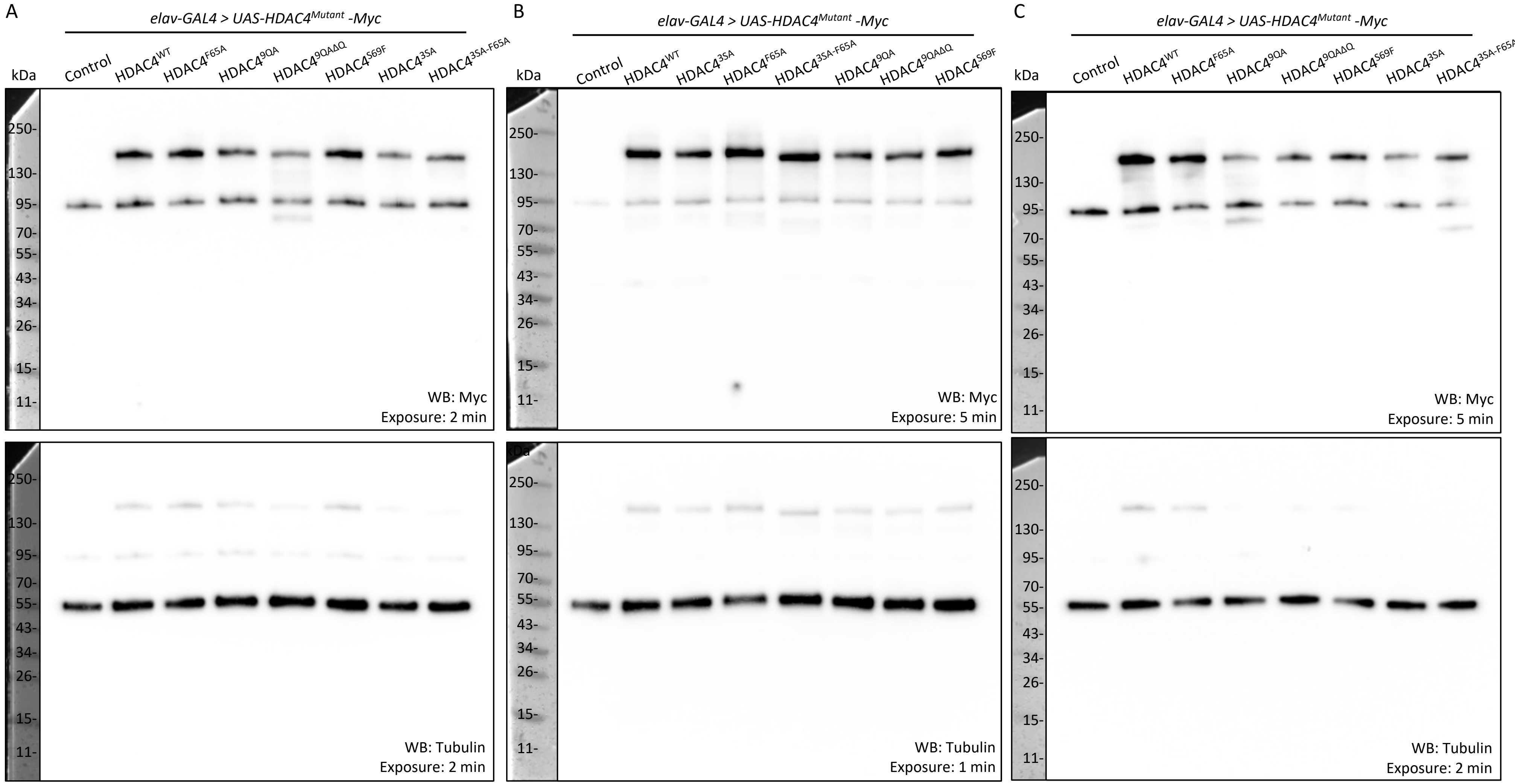


For the Myc detection, non-specific bands were observed at ~95 kDa in all samples including the negative control (*elav-GAL4/+*).

**Fig. 2**

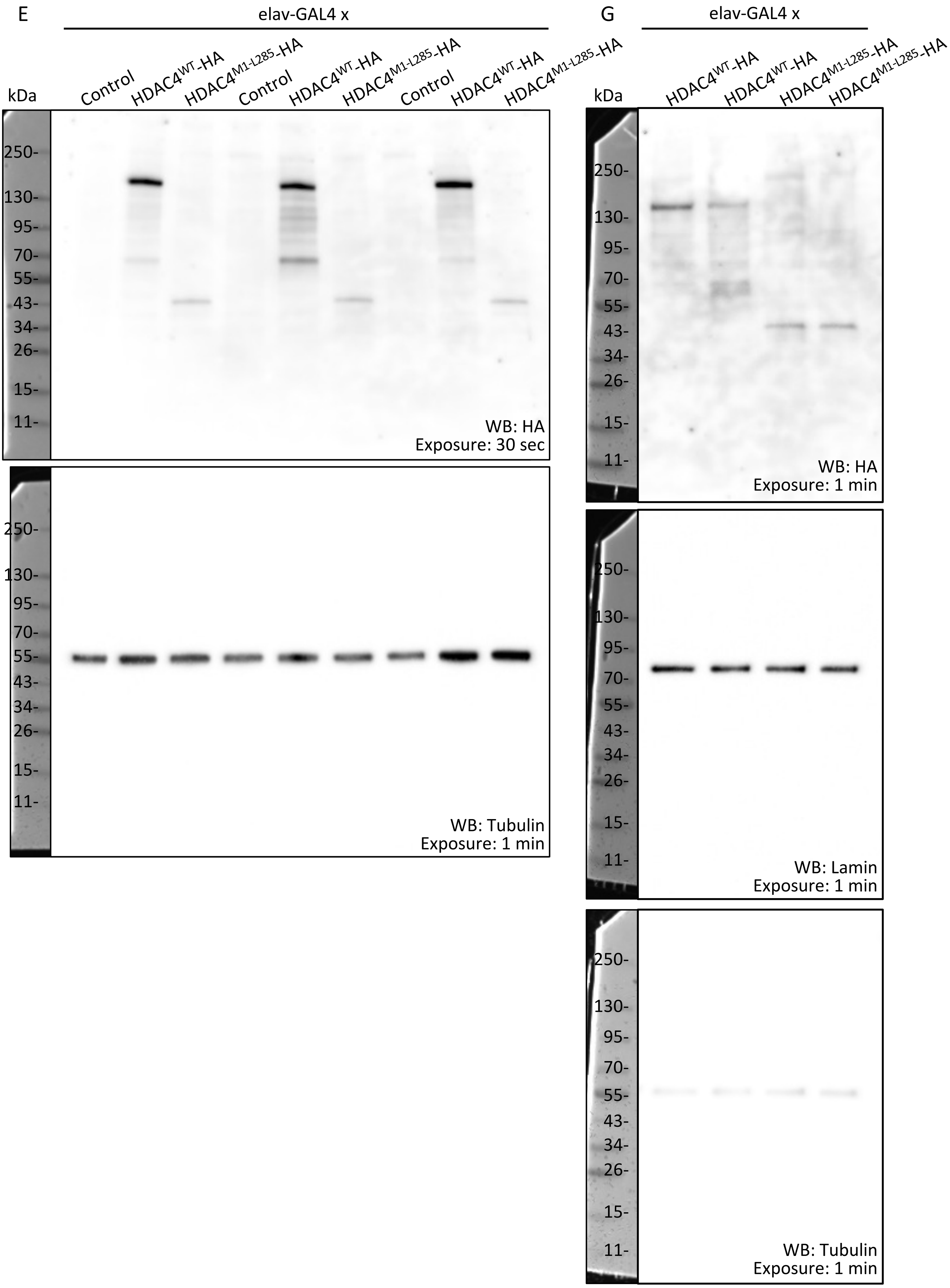


For the HA detection, non-specific bands were observed at ~250 kDa in all samples including the negative control (*elav-GAL4/+*).

**Fig. 7A**

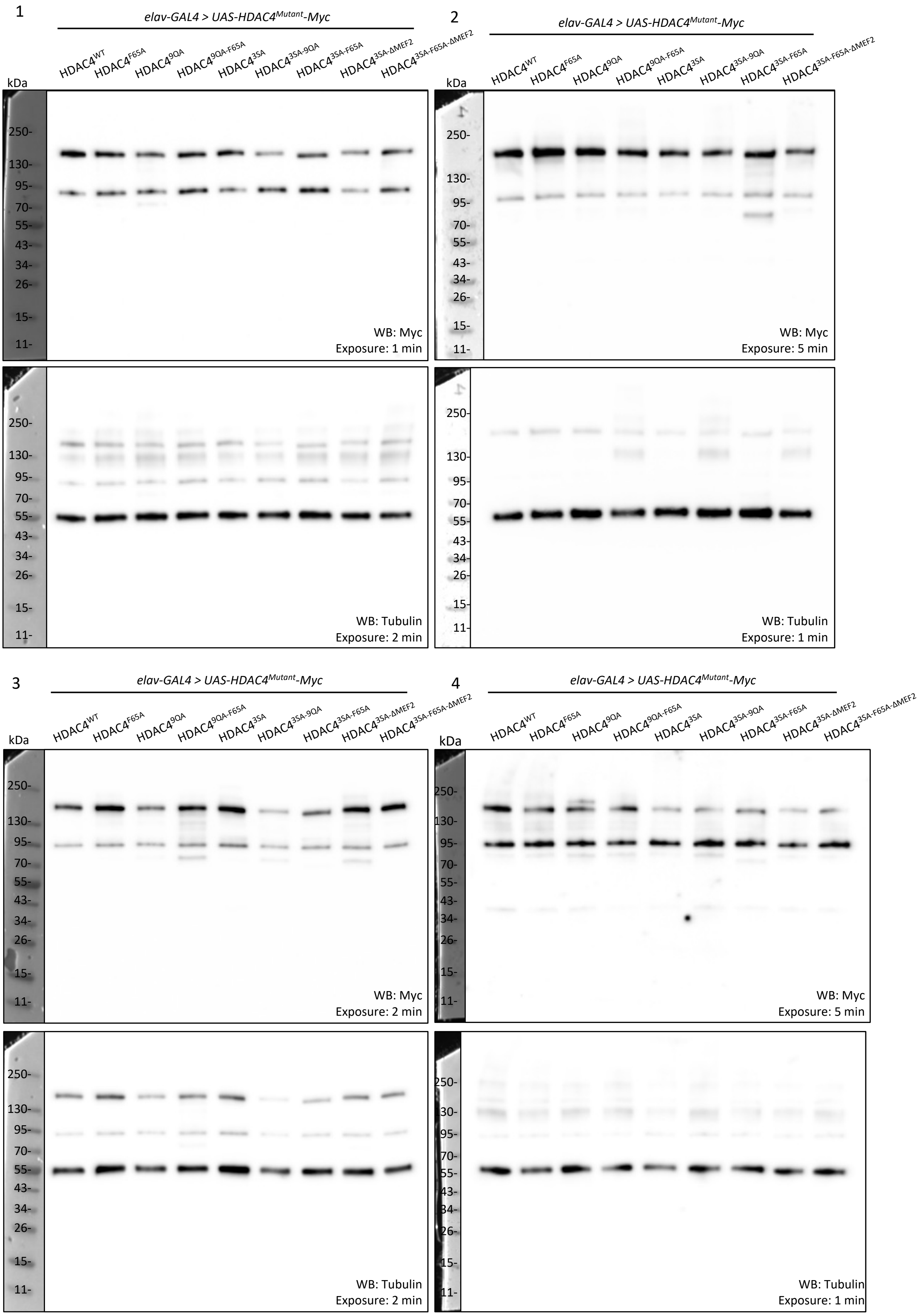


For the Myc detection, non-specific bands were observed at ~95 kDa in all samples as in Fig. 1H.

**Fig. 7 C-D**

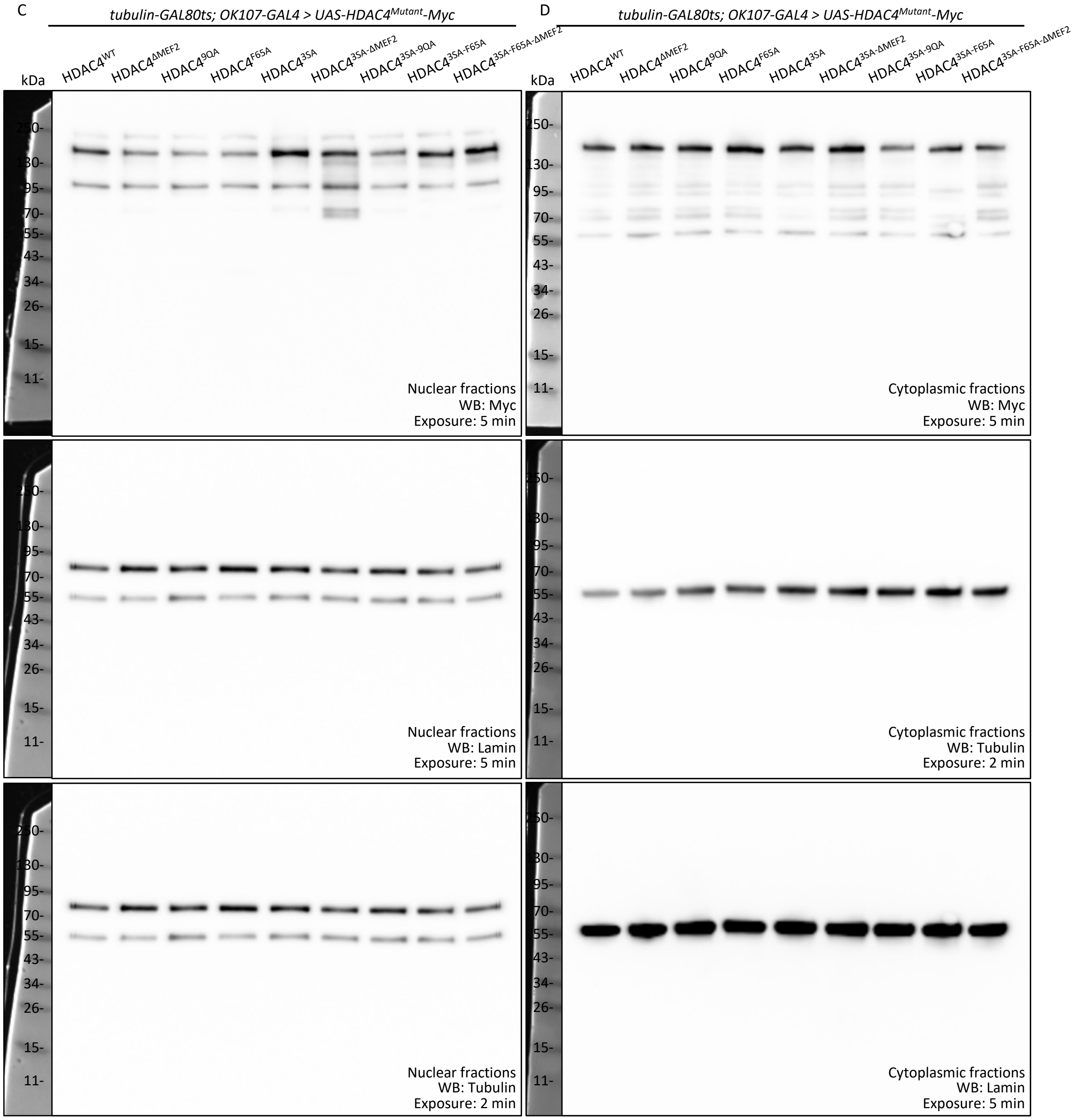


For the Myc detection, non-specific bands were observed at ~95 kDa in all samples as in Fig. 1H.

Blots were exposed for the same amount of time for each antibody detection and underwent the same adjustments in brightness and contrast between nuclear and cytoplasmic membranes.

Please note the same blot is shown for two different exposures for lamin and tubulin detections of nuclear fractions in (C). Lamin and tubulin were detected simultaneously.

**Fig. S2**

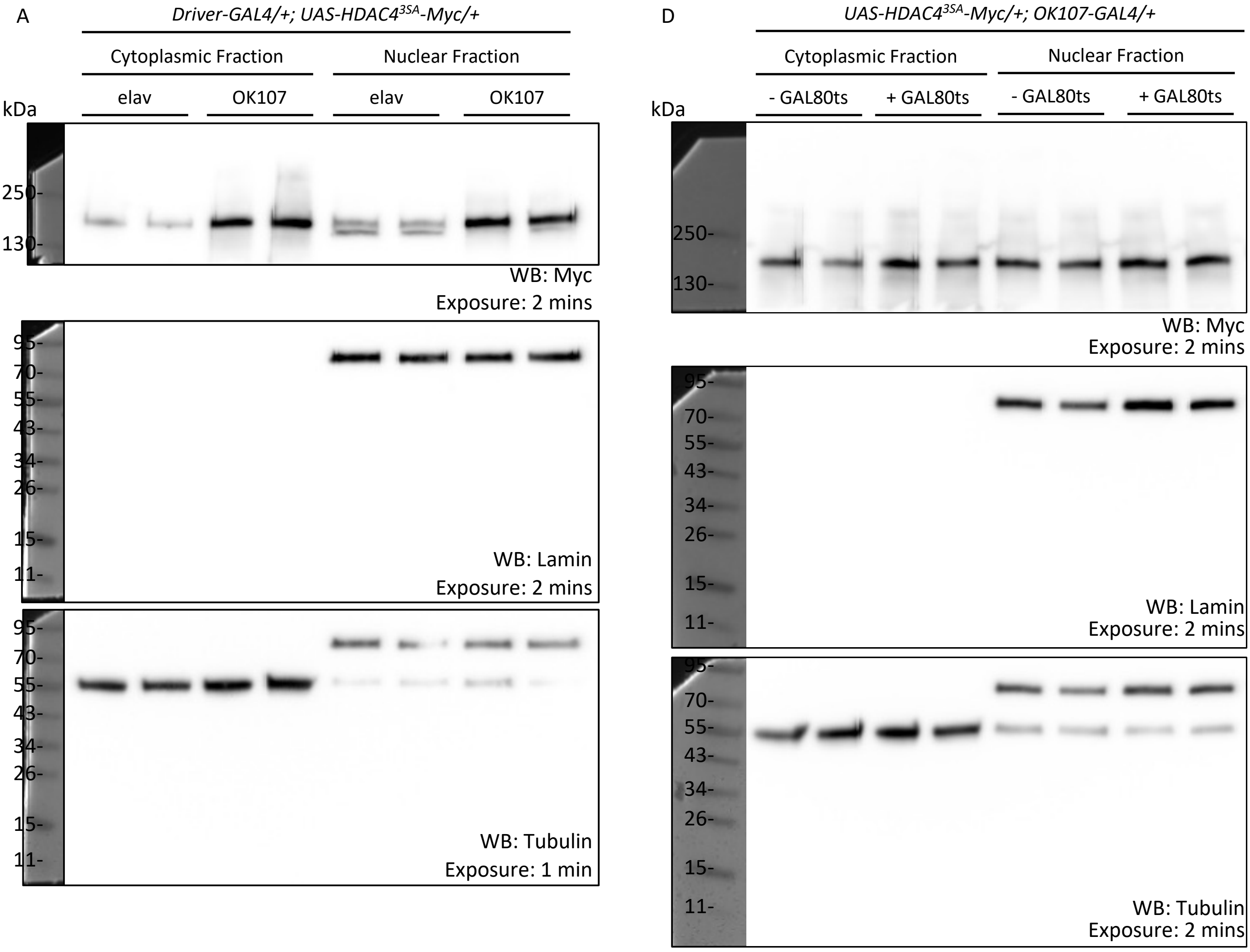


Blots were cut prior to detection. Bands at ~71 kDa in the tubulin detections are residual detection of lamin due to re-probing without stripping of the membrane.

**Fig. S7**

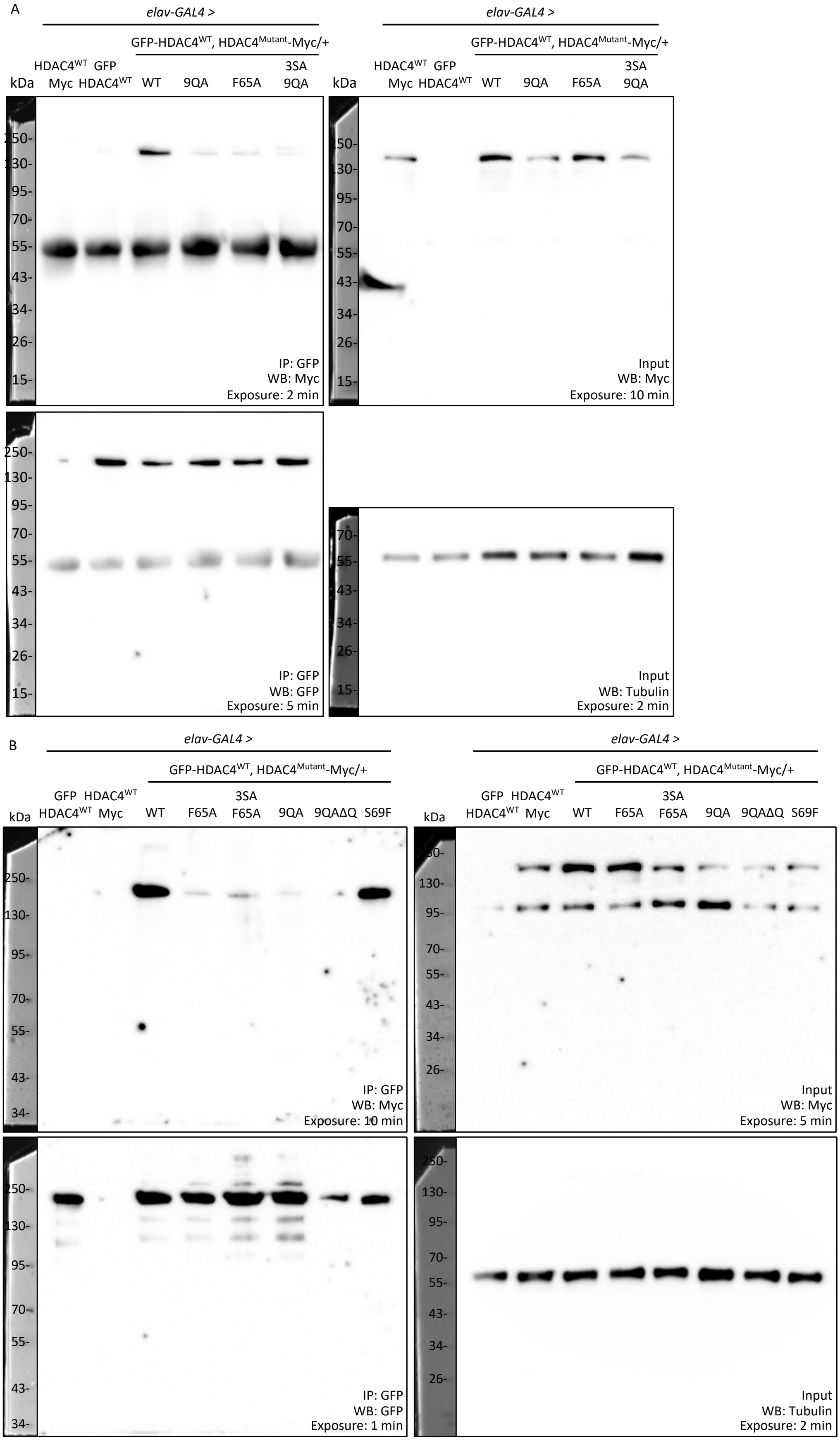


The band at ~50 kDa in the IP blots ((A), left) is IgG heavy chain. For the Myc detection in (B) (input, right), non-specific bands were observed at ~95 kDa in all samples as in Fig. 1H.
